# Multi-omics of barley Fusarium Head Blight converge on pathogen-triggered biosynthesis of aromatic amino acid derived chemical defense compounds

**DOI:** 10.1101/2025.09.08.674812

**Authors:** Sophia Hein, Christina E. Steidele, Felix Hoheneder, Sarah Brajkovic, Bernhard Kuster, Lisa Kurzweil, Timo D. Stark, Corinna Dawid, Ralph Hückelhoven

## Abstract

Fusarium Head Blight (FHB) is a devastating fungal disease of small grain cereals like wheat and barley, causing substantial yield and quality losses each year worldwide. FHB is caused by *Fusarium* species that produce mycotoxins such as deoxynivalenol (DON) that impairs protein biosynthesis. Although defense responses in barley to Fusarium infection have been described at the transcriptional level, it remains unclear to what extent these responses are translated into functional changes at the protein and metabolite levels.

In this study, we employed comprehensive transcriptomics, proteomics, and metabolomics to dissect the defense responses of barley heads during infection with *Fusarium culmorum*. Our integrated analyses revealed a set of significantly regulated gene-protein pairs linked to biosynthetic pathways that consistently correspond to upregulated defense-related metabolites. These include tryptophan-derived stress metabolites such as tryptamine and serotonin, as well as barley-specific hydroxycinnamylamides, including conjugates from the trypthophan metabolism, hordatines, and their biosynthetic precursors.

Integrating data across multiple omics layers identifies the upregulation of aromatic amino acid derived secondary metabolism as the most consistent barley response to FHB infection across diverse barley varieties that share barley-typical type II resistance to fungal spreading in the head rachis.

## INTRODUCTION

Fusarium Head Blight (FHB) is a devastating fungal disease affecting small grain cereals. The main causal agents for FHB in wheat and barley in Europe are *Fusarium graminearum* and *F. culmorum* (van der Lee et al. 2015). The soil-born pathogen infects the host plant at mid-anthesis through the open flower via ascospores or conidiospores that are dispersed through wind or rain splash (Moonjely et al. 2023). The hemibiotrophic fungi first grow biotrophically inside the host tissue before switching to a necrotrophic growth phase after approximately 48 hours. The necrotrophic growth phase leads to the visual symptoms of FHB like browning and withering of the infected spikelets (Wegulo et al. 2015). Fusaria produce a variety of mycotoxins which are mostly accumulating during the necrotrophic growth period, including Zearalenon, Nivalenol and Deoxynivalenol (DON) and its acetylated forms. DON is the primary mycotoxin of *F. graminearum* and *F. culmorum* (Desjardins et al. 1993). DON inhibits protein biosynthesis via binding to the peptidyl transferase center of eukaryotic ribosomes (Garreau de Loubresse et al. 2014). It is therefore also toxic for humans and animals, leading to serious health problems caused by contaminated grain (Rocha et al. 2005).

Given those economic and health implications of FHB, research into Fusarium resistance in small grain cereal crops like wheat and barley is important. In contrast to wheat, barley exhibits a natural type II resistance, preventing the spread of the fungal infection through the rachis, keeping it contained in the initially infected spikelets instead (Bethke et al. 2023; Boutigny et al. 2008). This natural resistance of barley makes it interesting for the study of immune and defense responses to *Fusarium* spp.. Previous research identified the UDP-glycosyltransferase HvUGT13248 as a key factor for type II resistance in barley. It detoxifies DON for the plant host by converting it to DON-3-Gucoside (Schweiger et al. 2010). Overexpression of the corresponding gene in Arabidopsis and wheat confers resistance (Li et al. 2017; Shin et al. 2012). In barley, overexpression of HvUGT13248 caused resistance to DON, and *HvUGT13248-*TILLING-mutants showed reduced FHB resistance compared to wild type barley plants (Bethke et al. 2023). However, it remains unclear which other factors contribute to pathogen defense against Fusarium in barley, including key metabolic pathways that might be involved in resistance to the pathogen.

Several studies have explored the transcript responses of barley to Fusarium infection. Upregulated genes included pathogenesis related (PR) genes, oxidative stress response genes, as well as genes involved in detoxification like glycosyltransferase and glutathione S-transferase genes (Boddu et al. 2006; Gardiner et al. 2010; Hameed et al. 2022; Hoheneder et al. 2023).

Proteomic studies on FHB in barley remain limited. Most existing research relies on 2D-gel based methods, limiting the depth of the analysis. Notably, proteomics approaches identified oxidative stress-related proteins, PR proteins, and enzymes involved in energy metabolism as responsive to FHB (Geddes et al. 2008; Yang et al. 2010). Artificial *F. culmorum* infection and DON treatment of barley affected proteins related to carbohydrate metabolism, amino acid metabolism, defense responses, redox regulation, and proteasome-mediated degradation (Kosová et al. 2017). However, no shotgun proteomics studies have been conducted in barley, highlighting a significant knowledge gap in this area. A number of metabolomics studies have identified key chemical groups and pathways linked to FHB resistance in barley. Several studies highlighted fatty acids, phenylpropanoids, terpenoids, amino acids, and hydroxycinnamic acids as important compounds associated with infection as well as resistance to FHB (Bollina et al. 2011; Chamarthi et al. 2014; Gauthier et al. 2015; Piasecka et al. 2022).

Although many studies investigated the complex responses of the host plant to Fusarium infection on single omics levels, understanding FHB in barley remains incomplete. Key questions about pathway involvement, protein expression, and the correlation between gene expression and metabolite production need a more comprehensive approach, combining transcriptomics, proteomics, and metabolomics, to give a more concise picture of host defense responses against Fusarium. Given the limited insights into the proteome, this study aims to overcome this gap by identifying key metabolic pathways involved in barley defense across multiple molecular levels. We seek to determine which pathways are consistently active in barley throughout transcriptomic, proteomic, and metabolomic responses, and how gene expression correlates with protein abundance and metabolite production. Such data provide valuable insights about which metabolic adaptations barley executes to limit fungal success during FHB pathogenesis. Here, we provide comprehensive multiomics data suggesting that barley responds to FHB infection by strong upregulation of aromatic amino acid derived chemical defense metabolism.

## RESULTS

### Comparison of transcripts and proteins identified in barley infected by *Fusarium culmorum*

To identify metabolic pathways in barley that contribute to the defense response against *F. culmorum*, one causal agent of FHB in barley, we investigated changes in gene expression, protein abundance and metabolite production after infection of barley head tissue. Four barley genotypes – Avalon, Barke, Morex, and Palmella Blue – were grown in controlled greenhouse conditions. The varieties share type II FHB resistance to fungal spreading through the rachis, but show differing levels of type I resistance to initial spikelet infection by FHB as previously observed in the field or greenhouse experiments (Hoheneder et al. 2022; Hoheneder et al. 2023). Avalon (Av) and Barke (Ba) are the more resistant and Palmella Blue (Pb) more susceptible variety, with Morex (Mo) showing moderate susceptibility. Around mid-flowering the heads of the plants were treated with either *F. culmorum* spore solution or water as Mock-treatment. The inoculated spikes were sampled 4 and 7 days after inoculation, respectively.

As a means to evaluate disease severity and infection success in the host, we quantified the contents of fungal DNA in relation to barley DNA through qPCR. The fungal DNA contents increased over the span of the sampling time points and differed between the barley varieties. Avalon and Barke exhibited comparable low infection levels. They showed at seven days post inoculation fungal DNA contents of 4.39 and 5.56 pg *Fcul* DNA/ng *Hv* DNA, respectively. Morex exhibited higher DNA contents at the same time with 9.63 pg *Fcul* DNA/ng *Hv* DNA and Palmella Blue contained the highest levels of fungal DNA (22.38 pg *Fcul* DNA/ng *Hv* DNA) (Table 1). Fungal DNA load thus reflects what has been reported earlier about quantitative resistance of these varieties (Hoheneder et al. 2022; Hoheneder et al. 2023). It however needs to be noted that due to differing flowering times, not all varieties could be inoculated simultaneously, but in two batches, with Avalon and Barke forming the first and Morex and Palmella Blue the second inoculation group. Comparability over all four varieties is therefore partially limited although data reflect genotype-typical disease progression.

**Table 1.** Fungal infection success as assessed by qPCR of fungal DNA content in barley spike tissue from four cultivars at four and seven days post inoculation (dpi) with *F. culmorum* spore solution.

| pg Fcul DNA/ng Hv DNA ± SD | | | | | | | |
| --- | --- | --- | --- | --- | --- | --- | --- |
| Avalon |  | Barke |  | Morex |  | Palmella Blue |  |
| 4 dpi | 7 dpi | 4 dpi | 7dpi | 4 dpi | 7 dpi | 4 dpi | 7 dpi |
| 2.04 ± 1.15 | 4.39 ± 1.29 | 2.39 ± 1.01 | 5.56 ± 1.58 | 0.95 ± 0.60 | 9.63 ± 5.06 | 1.23 ± 1.26 | 22.38 ± 10.32 |

To analyze gene expression after *F. culmorum* infection in the barley head tissue, we performed 3’-RNA sequencing. We mapped the RNA reads to the Morex-V3 reference genome (Mascher et al. 2021), evidencing RNA transcripts from 36920 gene models in all four varieties. The read counts of infected samples were then compared to the Mock-treated samples of the respective genotype and time point in order to find differentially expressed genes (DEGs). Each gene that shows a FDR-corrected P-value <0.05 in at least one comparison was considered a DEG. This led to a total number of 3085 DEGs in our dataset (Supplement Table D1.1). For 1205 of those DEGs, we additionally evidenced corresponding peptides in our untargeted proteomics analysis (see below; Figure 1 A). The number of differentially expressed genes found in each of the four tested cultivar differs, with Avalon, Palmella Blue and Barke containing around 1700 DEGs each, while in Morex 942 gene transcripts show differential expression after infection. All four cultivars share a set of 685 DEGs (Figure S1 A).

**Fig. 1.**
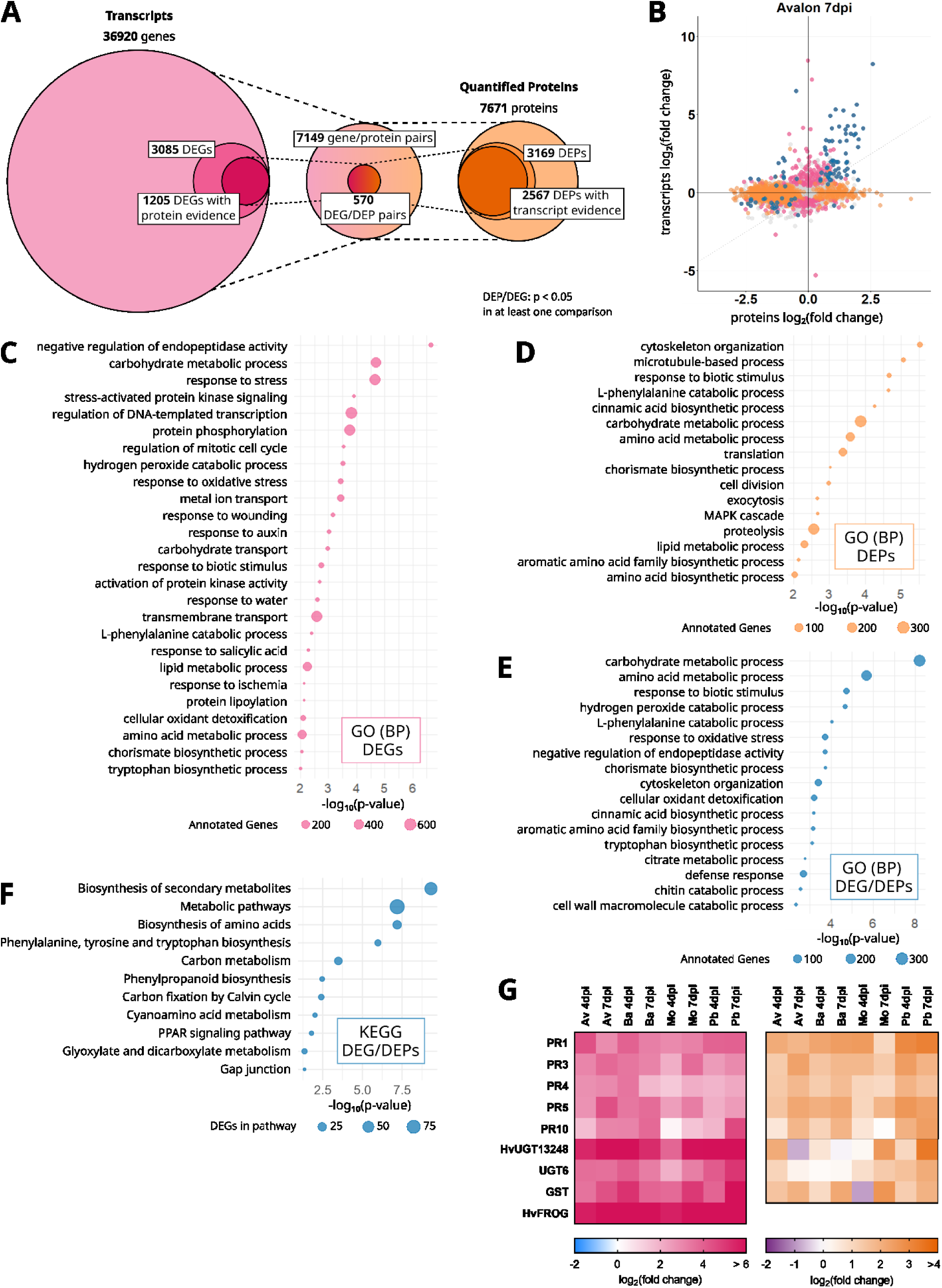
**A** Integrated analysis of gene transcript and protein abundances in barley spikes following *Fusarium culmorum* infection. Transcriptomic profiling was performed using 3′-RNA sequencing, with reads mapped to the Morex V3 reference genome, resulting in 36920 reliably detected gene models. Differential expression analysis (FDR-adjusted p < 0.05) between infected and mock-treated samples identified 3085 differentially expressed genes (DEGs). Proteomic analysis was conducted using untargeted, bottom-up proteomics with Tandem Mass Tag (TMT) labelling. A total of 7671 proteins were quantified, of which 3169 were classified as differentially abundant proteins (DEPs; p < 0.05). Transcript and protein datasets were integrated to identify 7149 gene/protein pairs, including 570 DEG/DEP pairs that showed significant regulation at both transcript and protein levels. These pairs represent prime candidates potentially involved in the barley defense response to *F. culmorum*. Transcript data are shown in pink and protein data in orange. **B** Comparison of fold changes of transcripts and corresponding proteins in barley cultivar Avalon 7 days post inoculation. Pink: Differentially expressed genes (DEGs). Orange: Differentially abundant proteins (DEPs). Blue: DEP/DEG pairs. **C** GO-term enrichment of 3085 DEGs. **D** GO-term enrichment of 3169 DEPs. **E:** GO-term enrichment of 570 DEP/DEGs. **F** KEGG pathway enrichment of 570 DEP/DEGs. **G** Differential expression of selected gene transcripts and proteins in barley spikes after inoculation with *F. culmorum* spore solution in comparison with mock-treated controls. Inoculation was performed around mid-anthesis, and samples were taken four and seven days post-inoculation (dpi). Infected samples were compared with mock-treated controls of the same cultivar and time-point. For each cultivar, treatment, and time point, four replicates were collected, each consisting of three pooled spikes. Heat-maps show the log₂(fold change) values for both gene transcripts and protein abundances. Colour keys for the heat-maps are provided below the panels. PR1: pathogenesis response gene 1 HORVU.MOREX.r3.5HG0444080; PR3: Chitinase HORVU.MOREX.r3.1HG0054950; PR4: pathogenesis response gene 4 HORVU.MOREX.r3.3HG0326310; PR5: Thaumatin-like protein HORVU.MOREX.r3.5HG0423100; PR10: pathogenesis response gene 10 HORVU.MOREX.r3.7HG0662380; *Hv*UGT13248: *Hordeum vulgare* UDP-glycosyltransferase 13248 HORVU.MOREX.r3.5HG0464880; UGT6: Glycosyltransferase HORVU.MOREX.r3.5HG0518060; GST: Glutathione-S-transferase HORVU.MOREX.r3.7HG0648120; *Hv*FROG: *Hordeum vulgare* Fusarium resistance orphan gene HORVU.MOREX.r3.4HG0345780.

To analyze changes in protein abundance after infection, we chose an untargeted bottom-up proteomics approach, including Tandem Mass-Tag (TMT) labelling to enable relative quantification of proteins. Peptide mapping and protein identification was carried out in MaxQuant using the Morex v3 genome-based protein prediction. This generated 6885 quantified protein groups, containing 7671 proteins (Supplement Table D1.2). We compared the normalized relative abundances of proteins between infected and Mock-treated samples of the respective genotypes and time points to calculate the fold changes in protein abundance. We considered all proteins with a p-value below 0.05 in at least one comparison as differentially abundant proteins (DEPs). The dataset includes 2813 DEP groups containing 3169 proteins (Figure 1 A, Supplement Table D1.3), with 1405 of those being Palmella Blue specific, followed by 591 and 345 Avalon and Barke specific DEPs, respectively. A subset of 83 DEPs are shared between all four genotypes (Figure S1 B). Among all quantified proteins and comparisons, tryptophan decarboxylase often showed the highest increase in abundance following *F. culmorum* infection. Several other strongly upregulated proteins included pathogenesis-related proteins, glycosyltransferases, and glutathione-S-transferases, all of which are commonly associated with plant defense responses to *Fusarium* species and DON.

The overlap of transcriptomics and proteomics datasets is comprised of 7149 gene/protein pairs for which we identified the transcript as well as quantified the corresponding protein. In order to find genes and proteins that are most reliably involved in the defense response against FHB, we concentrated on a subset of 570 genes and proteins that are both differentially expressed at transcript level as well as at protein level. Those we define as DEG/DEP pairs (Figure 1 A, Supplement Table D1.4 & D1.5).

Besides the stringent DEG/DEP pairs, a number of proteins show significant changes in abundance after infection while the corresponding gene transcript is not significantly regulated (Supplemental Table D1.3; Figure 1B, Supplemental Figure S2). Our dataset also includes 1997 DEPs without a significant change in gene expression levels in any of the barley varieties at any time. Of those, 553 are significantly upregulated in at least one cultivar at one time point. The remaining 1444 DEPs are downregulated in infected samples when compared to Mock-treated controls.

### GO-term and KEGG pathway Enrichment Analysis

To gain a better insight into what kind of biological functions are reflected within the sets of differentially regulated genes and proteins, we performed a GO-term enrichment analysis. For the 3085 DEGs this revealed a number of significantly enriched biological processes associated with stress and defense responses, including response to stress, hydrogen peroxide catabolic process, response to oxidative stress, response to wounding, response to auxin, response to biotic stimulus, response to salicylic acid, and cellular oxidant detoxification. In addition, several terms related to primary and secondary metabolism were enriched, such as L-phenylalanine catabolic process, lipid metabolic process, amino acid metabolic process, chorismate biosynthetic process, and tryptophan biosynthetic process. Additionally, enrichment of signaling and regulatory processes, such as stress-activated protein kinase signaling pathway, protein phosphorylation, and activation of protein kinase activity, was also significant (Figure 1C).

Analysis of the 3169 differentially expressed proteins (DEPs) revealed similar biological processes. GO terms related to primary and secondary metabolism were significantly enriched, including L-phenylalanine catabolic process, amino acid metabolic process, amino acid biosynthetic process, and cinnamic acid biosynthetic process. Defense-related responses like response to biotic stimulus were similarly present but DEPs more significantly showed translation, cytoskeleton and microtubule associated functions (Figure 1D).

Analysis of the 570 overlapping DEG/DEP pairs showed enrichment of defense responses and specialized secondary metabolism, notably cinnamic acid biosynthetic process, aromatic amino acid family biosynthetic process, tryptophan biosynthetic process, and amino acid metabolic process suggesting these pathways are enriched at both transcript and protein levels (Figure 1E).

In order to identify metabolic processes involved in the defense against Fusarium Head Blight in barley, we performed a *de novo* KEGG annotation of the Morex V3 genome, leading to the mapping of KEGG Orthology numbers (KO) to around 30% of amino acid sequences encoded in the reference genome (Supplement Table D1.6). With this we were able to perform a Pathway Mapping of the 570 DEG/DEP subset. The DEG/DEP pairs are significantly enriched for several pathways including Biosynthesis of amino acids, Phenylalanine, tyrosine and tryptophan biosynthesis, and Phenylpropanoid biosynthesis (Fig 1 F). The present pathways contain entries from 64 KEGG pathway modules, with three of those pathways being completely represented and nine only missing a single chain link (Supplement Table D1.7). Notable pathway modules include the shikimate pathway (M00022), tryptophan biosynthesis (M00023), phenylalanine biosynthesis (M00024), monolignol biosynthesis (M00039), flavanone biosynthesis (M00137) and melatonin biosynthesis (M00936).

### Differentially regulated genes and proteins in barley after *Fusarium culmorum* infection

The 570 stringently selected DEG/DEP pairs identified from 3’-RNA sequencing and bottom-up proteomics include a number of canonical pathogenesis-related genes (PR) and their respective proteins that are significantly upregulated after *Fusarium* infection. Several chitinase (PR3), Thaumatin-like protein (PR5) as well as PR1-genes show elevated transcript levels after infection nearly consistently throughout all barley varieties and time points with strong log_2_-fold changes of up to 4.20 (chitinase HORVU.MOREX.r3.3HG0277190; Ba 7 dpi), 10.95 (thaumatin-like protein HORVU.MOREX.r3.7HG0752040; Pb 7 dpi) and 7.00 (PR1 HORVU.MOREX.r3.7HG0668900; Pb 7 dpi), respectively (Figure 1G). For a majority of those gene transcripts, the corresponding proteins show a comparable regulation pattern with higher abundances after infection compared to the Mock-treatment (Figure 1G).

We furthermore find known Fusarium or DON response genes in the DEG/DEP subset, like UDP-glycosyltransferases, including a barley UGT6 ortholog (HORVU.MOREX.r3.5HG0518060) (He et al. 2020) and HvUGT13248 (HORVU.MOREX.r3.5HG0464880), which is known to glycosylate and detoxify DON (Schweiger et al. 2010). The transcript of this gene is strongly upregulated in all varieties with log_2_-fold changes ranging from 3.84 in Morex at 4 dpi to 7.75 in Barke at the same time point. This is, however, only partially translated into protein abundance, as in Avalon and Barke, the protein levels are slightly elevated after 4 days (log_2_-fold change: 2.12; 0.67), while they are even lowered compared to Mock at the later time point (log_2_-fold change: -0.78; -0.12). In Morex and Palmella Blue however, the abundance of HvUGT13248 increases after 7 days with log_2_-fold changes of 2.58 and 3.37, respectively, possibly reflecting the higher infection level of the fungus in those varieties (Figure 1G).

Among the strongly upregulated gene transcripts we found the barley ortholog of the Fusarium resistance orphan gene (HvFROG, HORVU.MOREX.r3.4HG0345780), which is associated with resistance to FHB in wheat (Perochon et al. 2015; Perochon et al. 2019). The respective log_2_-fold transcript changes range from 5.60 in Morex at 4 dpi to 11.43 in Palmella Blue at 7 dpi. The corresponding protein was not identified in our samples (Figure 1G). However, HvFROG is a small protein of predicted 142 amino acids and small proteins often evade detection in proteomics approaches.

### Metabolite abundance in barley after Fusarium culmorum infection

In an untargeted metabolomics measurement of the same plant material that we used for transcript and protein quantification, we detected over 14000 metabolic features. We could identify 53 unique metabolites either through reference substances measured on the same device (ID level 1), based on physicochemical properties and similarity to public libraries (ID level 2), or by similarity to known compounds of a compound class (ID-level 3). These include tryptophan and its derivatives, lipids and their oxidized fatty acid derivatives, hydroxycinnamic acids (HCAs), HCA-agmatines, as well as hordatines and their glucosides (Supplement Table D1.8) (Kurzweil et al. 2025). In order to identify additional hydroxycinnamic acid amides (HCAAs) in barley and gain information about their production under pathogen stress, we compared the spectral data to an *in silico* database of known and hypothetical HCAAs (Li et al. 2018). We focused on compounds with agmatine, serotonin and tryptamine as their amine moiety, because of the observed accumulation of hordatines, which contain agmatine as their amine component, as well as both serotonin and tryptamine after Fusarium infection. Methodological limitations can make it impossible to differentiate between different derivatives of a single HCAA like cis- or trans-isoforms, or compounds containing two or three hydroxycinnamic acid components. Therefore, several compounds are taken together under one annotation. We could identify several HCAAs or HCAA derivatives in barley, including caffeoyl-, cinnamoyl-, coumaroyl-, feruloyl-, and sinapoyl-agmatine; caffeoyl-, cinnamoyl-, feruloyl-, and sinapoyl-tryptamine; as well as caffeoyl-, cinnamoyl-, feruloyl-, and sinapoyl-serotonin (ID-level 3). Comparing Mock-treated and infected samples, caffeoyl-tryptamine, cinnamoyl-tryptamine, and cinnamoyl-serotonin significantly accumulated in barley after Fusarium infection, while the observed levels of sinapoyl-tryptamine, caffeoyl-serotonin, feruloyl-serotonin, and sinapoyl-serotonin are significantly decreased (Figure 2, Supplemental Figure S4 & S5).

**Fig. 2.**
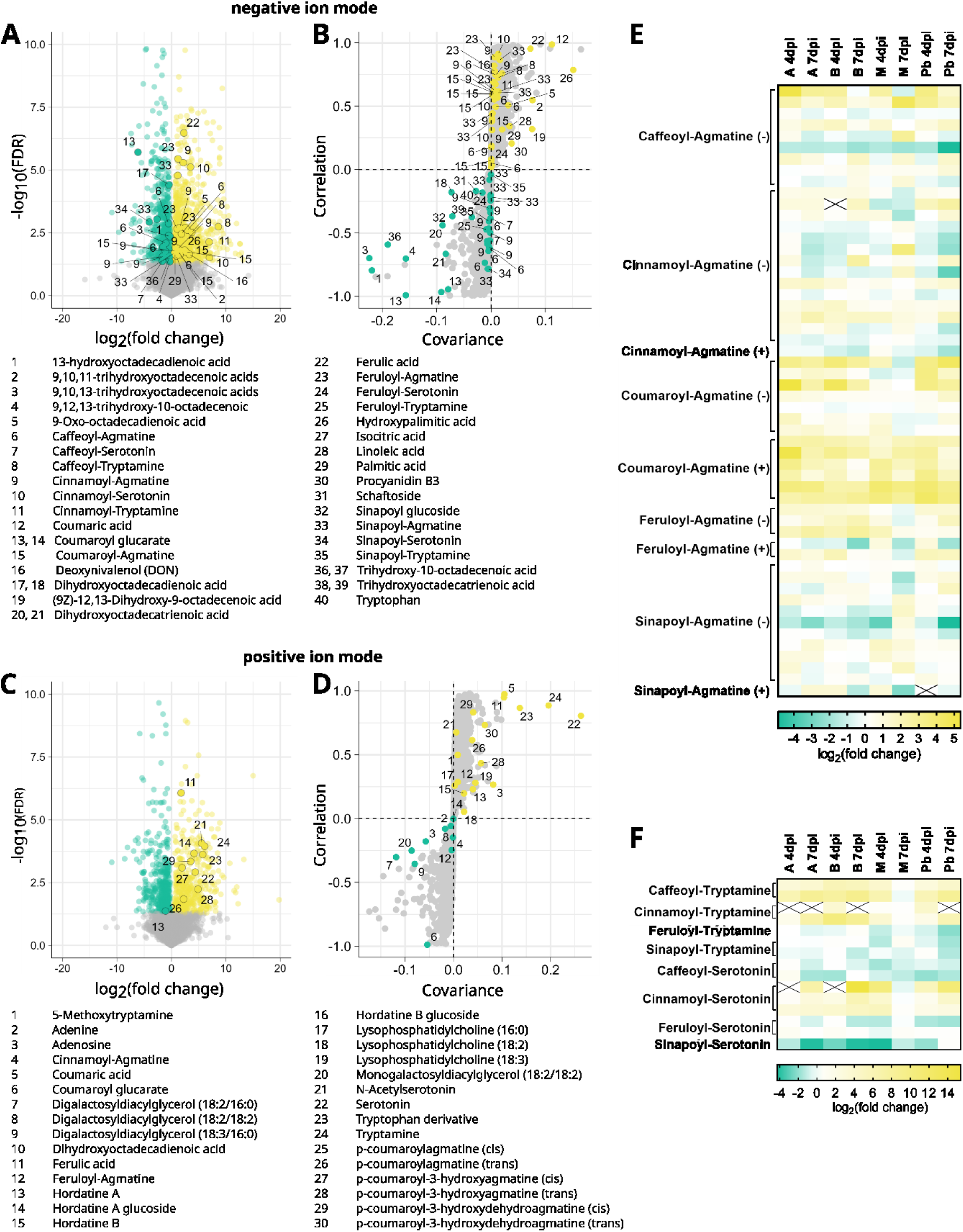
Differential abundance of metabolic features in barley cultivar Avalon at seven days post inoculation with *F. culmorum* spore solution in comparison with mock-treated controls. Inoculation was performed around mid-anthesis. Infected samples were compared with mock-treated controls of the same cultivar and time-point. For each cultivar, treatment, and time point, four replicates were collected, each consisting of three pooled spikes. Volcano plots show the log_2_(fold change) values in relation to the negative log_10_(FDR). S-plots show group differences as calculated via orthogonal partial least squares discriminant analysis (OPLS-DA). Metabolic features were measured via mass spectrometry in **A & B** negative ion mode and **C & D** positive ion mode. **E** Fold change in abundance of agmatine-containing hydroxycinnamic acid amides (HCAAs) in barley spikes after inoculation with *F. culmorum* spore solution in comparison with mock-treated controls. **F** Fold change in abundance of tryptophan-derived hydroxycinnamic acid amides (HCAAs) in barley spikes after inoculation with *F. culmorum* spore solution in comparison with mock-treated controls. Colour keys for the heat-maps are provided below the panels. Fields marked with a black X indicate a metabolite not being measured in the Mock-treated control samples, therefore preventing the calculation of a fold change.

Further analysis using orthogonal partial least squares discriminant analysis (OPLS-DA) revealed consistent infection-related metabolic patterns across all barley cultivars (Figure 2, Supplemental Figure S6 & S7). Lipid-derived metabolites, such as Lysophosphatidylcholines, and oxylipins, such as di- or tri-hydoxylated derivates of polyunsaturated fatty acids, were enriched or depleted depending on the individual substance suggesting a dynamic lipid metabolism in Fusarium-infected barley heads. Key marker compounds such as tryptophan-derived tryptamine and serotonin, p-coumaroylhydroxyagmatine (p-CHA), p-coumaric acid, and ferulic acid were upregulated in infected plants, while coumaroyl hexosides were depleted, suggesting that hydroxycinnamic acids may be further processed during infection (Figure 2, Supplemental Figure S4 & S5).

### Differentially regulated metabolic pathways after *Fusarium culmorum* infection

The KEGG pathway mapping revealed several connected pathway modules in the 570 DEG/DEP subset. The shikimate pathway, converting phosphoenolpyruvate and D-erythrose-4-phosphate to chorismate, is represented in the 570 DEG/DEP subset with several pathway genes, all showing upregulation at the transcript level after infection. In all genotypes except Barke, the log_2_-fold changes of most gene transcripts increased after 7 days in comparison to the earlier time point, as the infection progressed. This is partially translated into elevated protein levels. In the more resistant varieties Avalon and Barke, protein levels decreased at the later infection time compared to Mock potentially reflecting limited fungal progression (Figure 3).

**Fig. 3.**
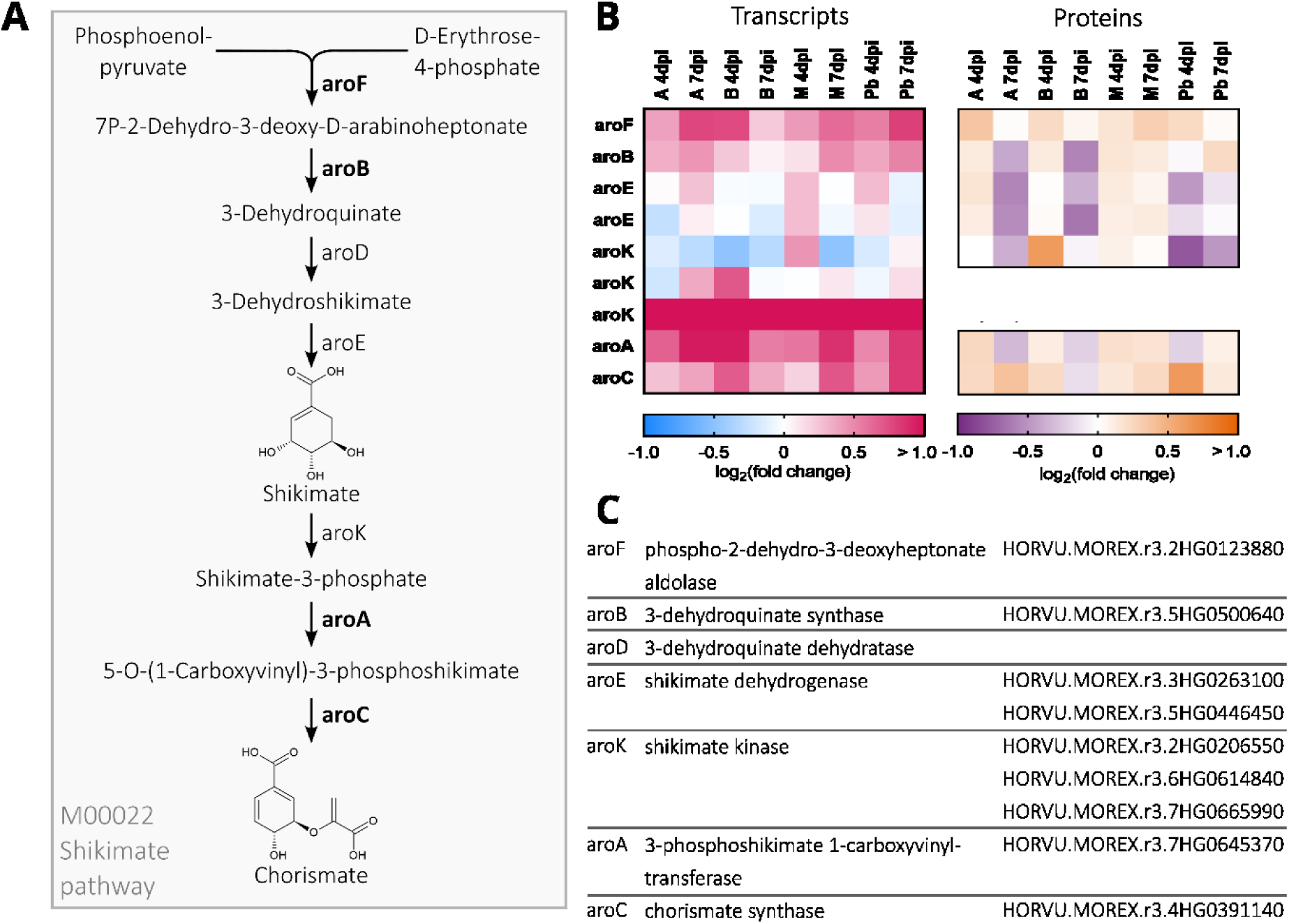
**A** Schematic overview of the shikimate pathway. Enzymes that are part of the 570 differentially expressed gene/protein (DEG/DEP) pairs are shown in **bold**. Fold changes are considered significant at a FDR-corrected P-value <0.05. **B** Differential expression of shikimate pathway gene transcripts and proteins in barley spikes after inoculation with *F. culmorum* spore solution in comparison with mock-treated controls. Inoculation was performed around mid-anthesis, and samples were taken four and seven days post-inoculation. Infected samples were compared with mock-treated controls of the same cultivar and time-point. For each cultivar, treatment, and time point, four replicates were collected, each consisting of three pooled spikes. Heat-maps show the log₂(fold change) values for both gene transcripts and protein abundances of shikimate pathway genes. Colour keys for the heat-maps are provided below the panels. **C** Pathway enzyme names and Morex v3 gene IDs.

Chorismate serves as a precursor for the biosynthesis of aromatic amino acids such as tryptophan. The KEGG pathway module for tryptophan biosynthesis is completely represented in the 570 DEP/DEG set, with all genes involved as well as their respective proteins being upregulated after infection across all varieties and time points. Tryptophan can be converted further to tryptamine and serotonin. Tryptophan decarboxylases (TDC), facilitating the conversion of tryptophan to tryptamine, are among the highest upregulated genes and proteins after infection in all varieties at both time points. While we can distinguish between seven individual TDC gene transcripts, on protein level they are all sorted into one protein group due to their peptide sequence similarity. We furthermore see a predicted tryptamine-5-hydroxylase (T5H) gene (HORVU.MOREX.r3.6HG0554760) upregulated at transcript level only. This also refers back to serotonin, the product of T5H enzyme activity, being among the metabolites most significantly and strongest enriched in infected samples compared to Mock in all varieties, except Morex at 4 days post-infection (Figure 2 & 4, Supplemental Figure S5).

Additionally, chorismate is the entry point for phenylalanine biosynthesis and its conversion into downstream metabolites like hydroxycinnamic acids (HCAs), HCA-CoA conjugates, aldehydes, and alcohols (Figure 5A). Several associated enzymes are found in the 570 DEG/DEP pairs. Most of the genes encoding those enzymes are upregulated, except shikimate O-hydroxycinnamoyltransferase (HCT) and one cinnamoyl-alcohol dehydrogenase (CAD) gene (Figure 5B). In most varieties and time points, this upregulation resulted in a moderate increase in protein levels at least at one time point. HCAs such as coumaric and ferulic acid were among the upregulated metabolites (Figure 5C).

**Fig. 4.**
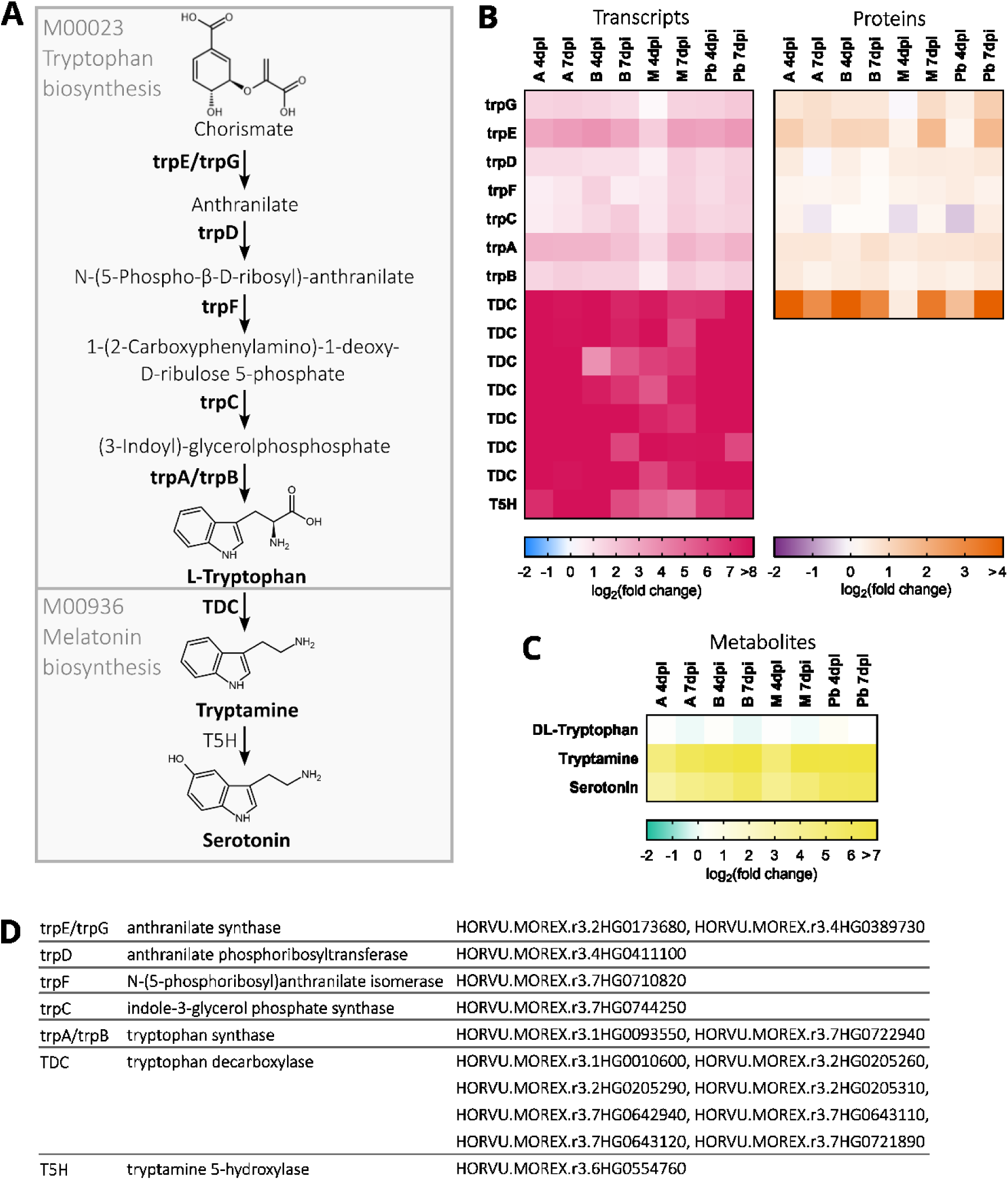
**A** Schematic overview of the tryptophan biosynthesis and melatonin biosynthesis pathways. Enzymes that are part of the 570 differentially expressed gene/protein (DEG/DEP) pairs are shown in **bold**. Fold changes are considered significant at a FDR-corrected P-value <0.05. **B** Differential expression of tryptophan and melatonin biosynthesis pathway gene transcripts and proteins in barley spikes after inoculation with *F. culmorum* spore solution in comparison with mock-treated controls. Heat-maps show the log₂(fold change) values for both gene transcripts and protein abundances of pathway genes. **C:** Fold change in abundance of metabolites in barley spikes after inoculation with *F. culmorum* spore solution in comparison with mock-treated controls. Inoculation was performed around mid-anthesis, and samples were taken four and seven days post-inoculation. Infected samples were compared with mock-treated controls of the same cultivar and time-point. For each cultivar, treatment, and time point, four replicates were collected, each consisting of three pooled spikes. Colour keys for the heat-maps are provided below the panels. **D** Pathway enzyme names and Morex v3 gene IDs.

**Fig. 5.**
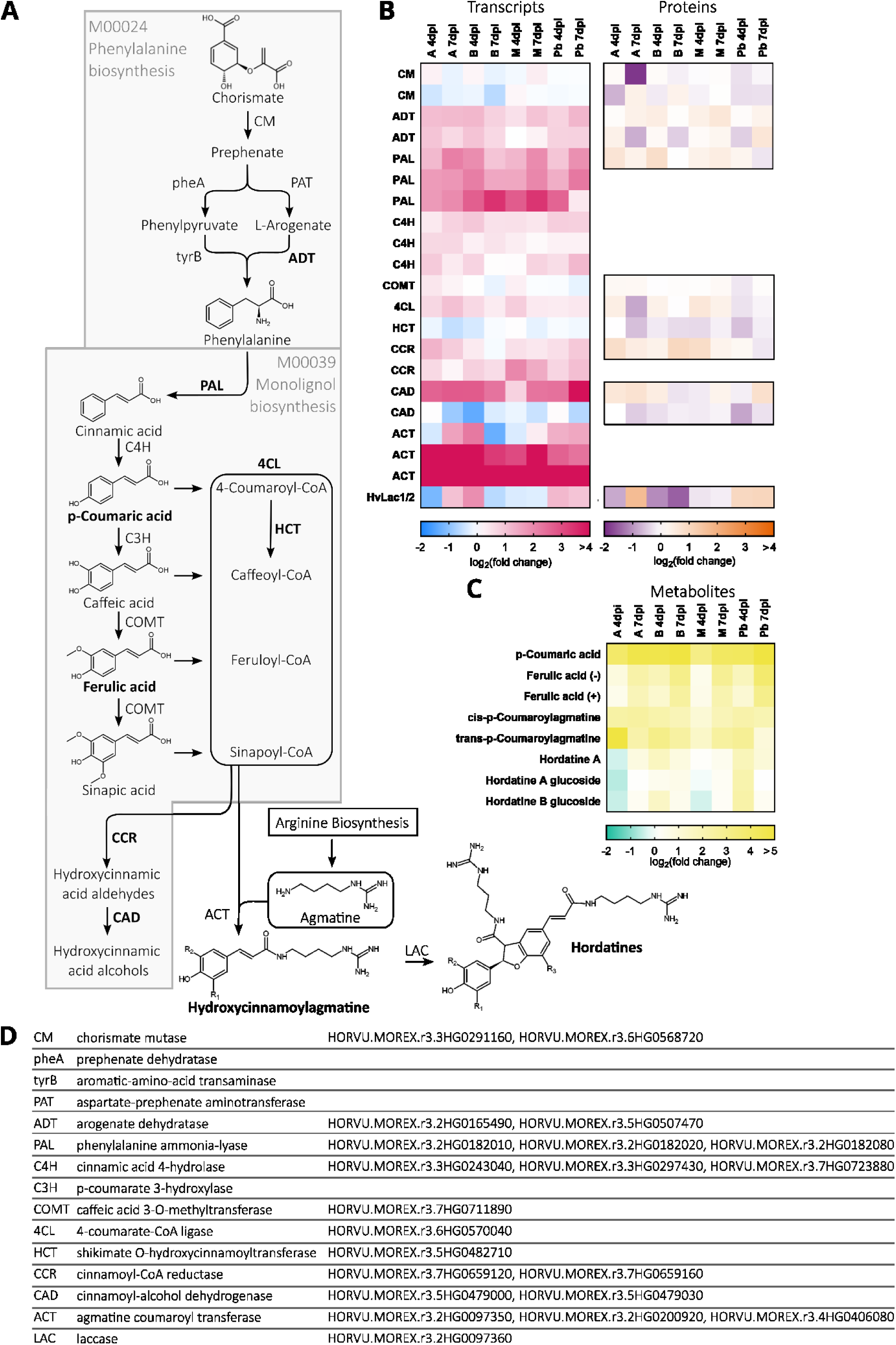
**A** Schematic overview of the phenylalanine biosynthesis and monolignol biosynthesis and hordatine biosynthesis pathways. Enzymes that are part of the 570 differentially expressed gene/protein (DEG/DEP) pairs are shown in **bold**. Fold changes are considered significant at a FDR-corrected P-value <0.05. **B** Differential expression of phenylalanine, monolignol and hordatine biosynthesis pathway gene transcripts and proteins in barley spikes after inoculation with *F. culmorum* spore solution in comparison with mock-treated controls. Heat-maps show the log₂(fold change) values for both gene transcripts and protein abundances of pathway genes. **C** Fold change in abundance of metabolites in barley spikes after inoculation with *F. culmorum* spore solution in comparison with mock-treated controls. Inoculation was performed around mid-anthesis, and samples were taken four and seven days post-inoculation. Infected samples were compared with mock-treated controls of the same cultivar and time-point. For each cultivar, treatment, and time point, four replicates were collected, each consisting of three pooled spikes. Colour keys for the heat-maps are provided below the panels. **D:** Pathway enzyme names and Morex v3 gene IDs.

The metabolites synthesized via the phenylalanine pathway contribute to the biosynthesis of HCA amides (HCAAs), metabolites known for anti-fungal and anti-microbial properties. In this context, it was interesting to see that two agmatine coumaroyltransferase-2 genes (HvACT; HORVU.MOREX.r3.2HG0200920, HORVU.MOREX.r3.4HG0406080) are highly upregulated on transcript level throughout all varieties at both sampled time points. However, we could not identify the ACT protein in our dataset and therefore cannot make any assumptions regarding its abundance. ACT synthesizes the coupling of agmatine with HCAs forming hydroxycinnamoylagmatines, HCAAs that can be further dimerized to form hordatines, barley-specific secondary metabolites (Figure 5A, B). Coumaroylagmatine as well as its dimer hordatine A and hordatine A glucoside show elevated metabolite levels after Fusarium-infection (Figure 5C).

The dimerization of hydroxycinnamoylagmatines to hordatines is potentially catalyzed by two Laccases HvLac1 and HvLac2 (Ube et al. 2023), which correspond to the gene HORVU.MOREX.r3.2HG0097360 in the Morex v3 reference. We evidenced the corresponding gene transcript and protein in barley head tissue, however not among the 570 DEG/DEP pairs.

In total, our multiomics analysis of the barley FHB response showed consistent upregulation of classical defense and detoxification responses and of pathways for aromatic amino acid-derived secondary metabolism.

## DISCUSSION

We combined three unbiased omics approaches that enabled an in-depth analysis of the defense response of barley against the hemibiotrophic pathogen *Fusarium culmorum*. Our dataset provides a comprehensive insight especially on transcript and protein level into the barley-Fusarium interaction on the host side. Single-omics studies might not show the complete picture of pathogen defense pathways or contain methodological bias. By contrast, the comparison between gene expression, protein abundance and metabolite levels from the same plant material identified pathways that are consistently upregulated at all molecular levels. Although plant varieties differed in basal FHB resistance, they all share barley-typical type II resistance that hinders spreading of the fungus through the rachis of the ear. We therefore hypothesize that commonly upregulated pathways might contribute to barley type II resistance against FHB.

### *Fusarium culmorum* infection in barley leads to gene-regulation and reshaping of the plant proteome

We could identify 36920 gene transcripts including 3085 DEGs after *F. culmorum* inoculation in at least one barley variety at 4 and/or 7 dpi. Additionally, we quantified 7671 barley proteins of which 3169 showed a significantly differential abundance during FHB infection. To focus on consistently regulated pathogen response pathways, we defined a subset of 570 DEG/DEP gene-protein pairs, showing both differential gene expression as well as differential protein abundance. This subset represents largely soluble proteins with successfully translated regulatory responses at the mRNA level and likely excludes gene regulation maintaining protein levels under stress or proteins insufficiently upregulated under the influence of *F. culmorum* or DON. This approach yielded identification of effectively employed metabolic pathways, which is well supported by our metabolomics results.

Previous omics studies mostly focused on single omics layers. The transcript responses of barley to *Fusarium* infection were investigated before, identifying upregulation of the tryptophan metabolic pathway, from chorismate to tryptamine synthesis (Boddu et al. 2006). A recent study on *F. culmorum* infection in barley identified an infection-related co-expression module including genes linked to defense responses, programmed cell death and mycotoxin detoxification, and a strongly overlapping gene set upregulated under FHB, including tryptophan decarboxylases, anthranilate synthase, laccases, and cinnamic acid metabolism genes (Hoheneder et al. 2023). Earlier studies identified overlapping sets of gene regulation in *Poacea* species in response to the related FHB pathogen *F. graminearum*. In barley defense related genes, oxidative stress, phenylpropanoid biosynthesis, and DON detoxification, were activated under *F. graminearum* infection as well as DON-exposure (Boddu et al. 2006; Boddu et al. 2007; Gardiner et al. 2010). Comparative transcriptomics in wheat revealed responses linked to hormone signaling, ROS burst, detoxification, and cell wall fortification (Buerstmayr et al. 2021; Wang et al. 2018). In a comparison of susceptible and moderately resistant wheat genotypes infected with *F. graminearum*, induced genes were enriched in aromatic compound metabolism, notably phenylalanine, as well as defense responses (Brauer et al. 2019).

Proteomic studies on FHB in barley or other Poaceae mainly relied on 2D-gel-electrophoresis, limiting analytical depth. Our shotgun-proteomics dataset therefore provides a more comprehensive view of translation of transcript-level defense responses. Former studies could mostly identify a limited number of differentially regulated proteins, including ones related to stress response, fungal defense, or oxidative stress (Kosová et al. 2017; Trümper et al. 2016; Yang et al. 2013). Gunnaiah et al. (2012) performed a shotgun proteomics analyses of QTL-mediated type II FHB resistance in wheat and reported similar activation of tryptophan metabolism and phenylpropanoid pathways leading to increased production of HCAAs and tryptophan-derived metabolites (Gunnaiah et al. 2012). Triticeae grasses may hence share a common FHB response in activation of those pathways. Individual metabolites like the hordatines, however, are often species or even genotype specific (Hamany Djande und Dubery 2024) and our data contain high numbers of unidentified metabolites enriched under FHB infection. Follow up structural analyses may reveal chemical defense compounds that correlate with spatiotemporal defense patterns and contribute to quantitative FHB resistance. Such substances further qualify as potential markers for resistance breeding (Baur et al. 2022; Dawid und Hille 2018; Hamany Djande und Dubery 2024).

In our dataset we identified 3169 differentially abundant proteins. The most highly upregulated ones include typical pathogen defense proteins like the pathogenesis related proteins, as well as known Fusarium responsive proteins with roles in mycotoxin detoxification like Glycosyltransferases and Glutathione-S-transferases. Those play an important role in the hosts response to Fusarium as well as barleys type II resistance to FHB (Bethke et al. 2023; Gardiner et al. 2010). The most consistently and strongly upregulated proteins were the tryptophan decarboxylases, key enzymes in serotonin biosynthesis. This is also reflected in tryptamine and serotonin being among the most upregulated metabolites in our samples after infection.

Another notable group comprises proteins with changes in abundance that cannot be explained by the regulation of their respective gene transcript. We identified 1997 of those proteins. Of these, 1444 are less abundant after infection, potentially due to the inhibition of translation through fungal toxins like Deoxynivalenol (DON). DON binds to the eukaryotic ribosome, inhibiting elongation (Garreau de Loubresse et al. 2014). Reduced abundance for those proteins could also be due to general stress-related protein turnover that is not compensated by gene up-regulation and altered cellular resource allocation. Additionally, 553 proteins show an increased abundance after infection in at least one cultivar at one time point, despite no significant transcript-level change. This might hint at post-translational modifications influencing protein stability and therefore leading to an accumulation of the protein. Deeper analysis of this subset of proteins could provide insight into post-translational modifications in response to *Fusarium* infection. This would require additional proteomics experiments such as phospho-enrichment or building a tryptic peptide library including possible modifications. As this study aimed to investigate core FHB responses, we did not further analyze these discordant cases, which could be addressed in future work.

The DEG/DEP subset likely does not capture the full spectrum of induced defense responses, as methodology limits the depth of the analysis, especially for proteomics. The transcript dataset contains 36920 gene transcripts, while we quantified 7671 barley proteins, meaning that proteins not detected by shotgun proteomics are absent from our dataset. A notable example is the Fusarium resistance orphan gene (HvFROG, HORVU.MOREX.r3.4HG0345780), associated with FHB resistance in wheat (Perochon et al. 2015; Perochon et al. 2019). Although the transcript is highly upregulated, the corresponding protein is not detected, suggesting that small or low-abundance proteins may escape untargeted LC-MS workflows. Targeted approaches may therefore be required to capture such proteins when they are of particular interest. However, as our primary aim was to identify broader metabolic trends, a shotgun strategy remained the most appropriate choice for this study.

### Differences between barley cultivars do not reflect disease susceptibility

Differences between the tested cultivars are evident in fungal DNA levels, reflecting variation in susceptibility. Morex and Palmella Blue showed the highest fungal DNA contents after 7 days, while Avalon and Barke had considerably lower levels, indicating a stronger resistance to Fusarium infection (Table 1). It is important to note that two separate infection events occurred, which limits direct comparison between samples. However, the core response of diverse barley genotypes to FHB infection rather than genotype comparisons are in the focus of this study, so this limitation is not of major concern.

Several barley varieties were included to identify reliable pathways that play a role in FHB across a diverse panel of cultivars. Differences in flowering time reflect physiological differences and cannot be avoided without introducing further batch effects e.g. by different sowing dates. Inoculum batches were prepared with consistent spore concentrations using plates of similar growth times and counting the spore concentration for each batch. Growth conditions in the greenhouse were kept stable between the inoculation events to ensure a comparable infection success and initial fungal DNA load was comparable at 4 days after inoculation before differences in susceptibility appeared. We therefore assume that differences in quantified fungal DNA reflect different susceptibility of the chosen cultivars rather than inoculum or environmental variation. Differences in fungal load are not always apparent from the observed fold changes of gene transcripts and protein abundance. In many cases, fold changes in gene expression appear quite similar across all cultivars. This suggests that quantitively type I resistant and susceptible cultivars respond similarly to fungal infection. However, resistant cultivars reach the same upregulation while confronted with less fungal load. Data might also contain a dilution effect, as our samples contained the whole head, with infected and healthy tissues alike. This stems from spray inoculating the entire spike rather than single spikelet inoculation. This approach mimics natural infection more closely where spores land on spike tissues and grow into the open spikelet. However, this could lead to an underestimation of the defense response in resistant cultivars.

Some pathogen responses went down again at 7 dpi in the more resistant cultivars, which could further indicate that the fungal development was already restricted at this time in the more resistant cultivars. Previous studies showed delayed defense responses in susceptible cultivars. A recent shotgun-proteomics study investigated the role of Fusarium mycotoxins on the wheat proteome. Treatment of resistant and susceptible wheat varieties with the acetylated DON-derivative 15-ADON led to substantial remodeling of the proteome in response to toxin treatment. Toxin-specific protective responses like the production of glutathione were delayed in the susceptible compared to the resistant cultivar (Buchanan et al. 2025). Similarly, *Zymoseptoria tritici* leads to the accumulation of HCAAs in infected wheat with susceptible cultivars showing a delayed accumulation of the compounds compared to resistant cultivars (Seybold et al., 2020). Future work should thus include approaches that reach a higher spatial and temporal resolution of the molecular responses.

### *Fusarium culmorum* infection leads to metabolite accumulation in barley

We quantified 14218 metabolic features, but discuss here mainly substances that have been evidenced by the help of authentic standards (level 1) or by similarity in physicochemical properties and published MS spectra (level 2) (Supplement Table D1.8). Among the 40 metabolites identified with high certainty, we can find key compounds that showed antifungal or defense properties in previous studies. Serotonin is upregulated in all tested genotypes and was previously shown to play a role in the immune response of rice to *Magnaporthe grisea* and *Bipolaris* (Fujiwara et al. 2010; Hayashi et al. 2016; Ishihara et al. 2008; Kurzweil et al. 2025). It plays a multi-faceted role in plant development and stress response, promoting seed germination as well as regulating plant growth via crosstalk with plant hormones like auxin. Serotonin is also linked to abiotic and biotic stress tolerance (Mishra und Sarkar 2023; Sun et al. 2025). In *Poacea*, it accumulates after pathogen attack, contributes to mechanical barriers against invading pathogens, and reduces biotic stress in rice by producing halo lesions against *Magnaporthe oryzae* (Hayashi et al. 2016; Ishihara et al. 2008). However, the role in pathogen defense depends on the plant species and remains mechanistically unclear (Erland et al. 2016). Additionally, serotonin and tryptamine can act as substrates for hydroxycinnamoyltransferases to generate bioactive HCAAs (Moghe et al. 2023). Hydroxycinnamic acids (HCAs), namely p-coumaric acid, ferulic acid, and cinnamic acid, were found to accumulate in wheat and barley upon infection and inhibit growth and conidia germination of *F. graminearum* (Fajardo et al. 2025; Ube et al. 2019b). Treatment of *F. oxysporum* f. sp. *niveum* with high concentrations of ferulic acid *in vitro* had a negative effect on fungal growth and inhibited conidia germination (Wu et al., 2010). Hydroxycinnamic acid amides (HCAAs) were shown to exhibit similar effects. HCAAs are believed to have two major protective functions, antimicrobial activity against bacteria and fungi and reinforcement of the cell wall (Roumani et al., 2020). Feruloylagmatine inhibited spore germination of *F. pseudograminearum in vitro* in a previous study (Carere et al. 2018), and HCAAs like cis-p-coumaroylagmatine and feruloylagmatine increased cell wall thickness in wheat infected with *F. graminearum* (Gunnaiah et al. 2012). HCAAs can be incorporated into the cell wall, crosslinking between structural polymers like polysaccharides via ester bonds and increasing rigidity (Zeiss et al., 2020).

Among HCAAs, hordatines, dimerized cinnamoylagmatines, are of particular interest as they are specific to cultivated and wild barley (Smith and Best 1978; Stoessl and Unwin 1970). Hordatines and their precursors p-coumaroylagmatines were shown to accumulate after infection of barley with powdery mildew (Röpenack et al. 1998; Smith und Best 1978). Hordatines further accumulate in barley exposed to powdery mildew resistance-inducing volatile organic compounds released from wounded or infected barley plants (Laupheimer et al. 2023; Laupheimer et al. 2024). Hordatines were tested for their antifungal effects in spore germination assays. Hordatine A inhibits spore germination of several fungi including *F. oxysporum* at concentrations as low as 10^-5^ M in liquid culture, though effects on agar plates and mycelial growth are limited (Stoessl and Unwin, 1970). HCAA accumulation after pathogen stress occurs across many pathosystems (Macoy et al. 2015; Moghe et al. 2023). Similar to our findings, serotonin and *p*-coumaroyl-hydroxydehydroagmatine were previously shown to be significantly elevated in *Ramularia collo-cygni*-infected barley leaves, further supporting their role in pathogen defense (Lemcke et al. 2021).

While hordatines and hydroxycinnamoyl-agmatines are already well described compound classes in barley, data on the prevalence of other HCAAs is lacking (Hamany Djande et al. 2022; Hamany Djande und Dubery 2024). Using supervised annotation based on an *in silico* database, we identified several HCAA derivatives, including caffeoyl-, cinnamoyl-, feruloyl-, and sinapoyl-agmatine, -tryptamine, and -serotonin (Li et al. 2018). We carefully classified these compounds as ID-level 3 due to methodological limitations that make it impossible to distinguish between derivatives. However, accurate mass, fragments, retention time prediction, and spectral matching increase confidence to surpass ID-level 3. While coumaroyl- and feruloyl-agmatine and their downstream products accumulate after Fusarium infection, as well as derivatives of caffeoyl- and cinnamoyl-tryptamine, and cinnamoyl-serotonin, other HCAAs like sinapoyl-tryptamine, caffeoyl-, feruloyl-, and sinapoyl-serotonin show negative fold changes when comparing infected to Mock-treated samples. Depletion of those compounds could be related to incorporation of specific HCAAs into the cell wall, as was shown previously for cis-p-coumaroylagmatine and feruloylagmatine in wheat infected with *F. graminearum* (Gunnaiah et al. 2012). Incorporation into the cell wall makes metabolic compounds inaccessible for methanolic extraction and subsequent LC-MS measurement. As cell wall fortification is a plausible mode of action for hordatines and other HCAAs this makes the assessment of the production of these compounds difficult. In order to validate the incorporation of HCAAs into the cell wall during infection, additional experiments would be needed.

Serotonin- and tryptamine-containing HCAAs represent the chemical convergence of the two infection-responsive metabolic pathways identified in this study. The presence and accumulation of those compounds further highlights the coordinated activation of both the tryptophan and hydroxycinnamic acid biosynthesis pathways. Some of the HCAAs we could identify in barley have also been previously described in other grasses. In rice leaves, tyramine and feruloyl-agmatine are induced upon infection with *Cochliobolus miyabeanus* or *Xanthomonas oryzae*, and strong accumulation of tyramine-derived HCAAs has also been observed during infection by *X. oryzae* and *Magnaporthe oryzae* (Xue et al. 2025).

### *Fusarium culmorum* infection in barley leads to the upregulation of metabolic pathways converging on aromatic amino acid derived compounds

The *de novo* KEGG annotation of the barley genome and pathway mapping of the 570 DEG/DEP subset identified 64 regulated KEGG pathway modules (Supplement Table D1.7). Comparison between the pathway modules and the metabolites increasing after infection identified 6 KEGG pathway modules involved in the biosynthesis of HCAAs and their precursors, whose end products include barley-specific hordatines accumulating during FHB. Those pathways align closely with enriched GO-terms, linking to primary and secondary metabolism. Those include aromatic amino acid biosynthetic process, cinnamic acid biosynthetic process, and chorismate biosynthetic process. Together, these findings underline the significance of this pathway for the barley defense response to *Fusarium* infection.

Serotonin is among the strongest upregulated metabolites, placing tryptophan metabolism centrally in barley’s FHB response. Tryptophan decarboxylase protein regulation reached stronger increase levels than canonical PR-proteins. Additionally, we found HORVU.MOREX.r3.6HG0554760 strongly upregulated (Figure 4B), which is the putative barley ortholog of the rice tryptamine-5-hydroxylase (86% protein identity, 92% protein similarity) generating serotonin (Ameen et al. 2021; Fujiwara et al. 2010). This function was recently confirmed by heterologous expression experiments (Christensen et al. 2025). The same study showed strong activation of the tryptophan metabolism in barley leaves after infection with the hemibiotrophic pathogen *Pyrenophora teres* f*. teres*. They specifically observed an accumulation of serotonin and two hydroxylated tryptamine isomers, including 2-oxo-tryptamine, at the site of infection. Together with our findings on another pathogen infecting another plant organ, this puts tryptophan-derived secondary metabolites at a potentially central position in barley chemical defense.

Agmatine coumaroyltransferases (ACTs) as key enzymes for the formation of HCAAs were linked to resistance in previous studies and showed significant transcriptional upregulation in our dataset. In *Brachypodium distachyon*, loss-of-function mutants of ***Bd*ACT2a** exhibited reduced accumulation of feruloylagmatine and increased susceptibility to *F. pseudograminearum* (Carere et al. 2018). Similarly, in wheat, ***Ta*ACT** was identified within a quantitative trait locus (QTL) associated with FHB resistance. Virus-induced gene silencing of *Ta*ACT enhanced susceptibility to *F. graminearum* and lowered levels of p-coumaroylagmatine (Kage et al. 2017).

Barley naturally exhibits a higher type II resistance to FHB compared to other small grain cereals like wheat. Detoxification mechanisms like DON glucosylation via barley *Hv*UGT13248 were previously shown to contribute to type II resistance (Bethke et al. 2023). However, it is not known whether additional host defense responses are involved in type II resistance of barley. All cultivars used here share, despite strong differences in type I resistance to initial infection, full type II FHB resistance. This raises the possibility that core chemical defense factors may contribute to barley’s response to FHB. In this context, hordatines or other aromatic amino acid–derived secondary metabolites emerge as potential candidates, but their functional relevance remains to be experimentally validated. Literature further suggests the formation of HCAA from HCA and tryptophan-derived amines such as tryptamine or serotonin leading to so-called Triticamides in *Triticeae* (Ube et al. 2019a). Since our data shows that metabolic reprogramming of barley for the FHB response converges on HCA- and tryptophan-metabolism, we suggest future investigations of the chemical diversity of predescribed and potentially new HCAAs that emerge as conjugates from those pathways. A number of additional Fusarium-responsive laccases (HORVU.MOREX.r3.1HG0008790, HORVU.MOREX.r3.1HG0020450, HORVU.MOREX.r3.3HG0313580, HORVU.MOREX.r3.4HG0388460, HORVU.MOREX.r3.6HG0614790, HORVU.MOREX.r3.7HG0637080), other than hordatine-forming Lac1/2 (Ube et al. 2023), in our dataset (Supplement Table D1.4 & D1.5) leaves further room for speculations on the formation of HCAA dimers or oligomers beyond known hordatines.

## CONCLUSION

This study provides a comprehensive multiomics analysis of Fusarium Head Blight disease and chemical defense responses in barley. This allows for insight into which transcriptomic responses are actually translated into metabolic read-outs and can help to identify candidate defense compounds that play a role against fungal attack. We see strong reactions in PR-proteins and known Fusarium responsive genes as well as metabolic pathways producing aromatic amino acid derived chemical defense compounds like serotonin, HCAAs and hordatines, with three unbiased omics approaches all converging on secondary metabolite pathways around those metabolites. Among those, hordatines are prime examples for Fusarium-induced HCAAs that are specific for barley as a host organism, hinting at possible factors for the higher type II resistance of barley to FHB in comparison with wheat. Future genetic analysis of HCAAs metabolism will shed light on its impact on FHB resistance of barley.

## LIST OF ABBREVIATIONS

ACT: Agmatine coumaroyltransferase
Av/A: Avalon
Ba/B: Barke
bRP: Basic reversed-phase (fractionation)
DEG: Differentially expressed gene
DEP: Differentially abundant protein
DON: Deoxynivalenol
dpi: days post inoculation
FDR: False discovery rate
FHB: Fusarium head blight
GO: Gene Ontology
GST: Glutathione-S-transferase
HCA: Hydroxycinnamic acid
HCAA: Hydroxycinnamic acid amide
KEGG: Kyoto Encyclopedia of Genes and Genomes
KO: KEGG orthology
LC-MS/MS: Liquid chromatography coupled to tandem mass spectrometry
Mo/M: Morex
MS: Mass spectrometry
OPLS-DA: Orthogonal partial least squares discriminant analysis
Pb: Palmella Blue
PCA: Principal component analysis
PR: Pathogenesis-related
T5H: Tryptamine-5-hydroxylase
TDC: Tryptophan decarboxylase
TMT: Tandem mass tag
UGT: UDP-glycosyltransferase
UHPLC-TOF-MS: Ultra-high-performance liquid chromatography time-of-flight mass spectrometry

## MATERIALS AND METHODS

### Infection experiments

Greenhouse experiments consisted of four barley genotypes of differing susceptibility to Fusarium Head Blight, namely Barke, Avalon, Morex and Palmella Blue (Hoheneder et al. 2022). The plants were grown under controlled conditions in a greenhouse cabin on flooding tables at the Plant Technology Center of the Technical University of Munich. 3-litre pots with a peat substrate (Einheitserde C700, Stender, Germany) were used to grow five seeds each at a temperature of 18/16 °C day/night, with additional lighting for 16 hours per day and a relative air humidity of 60%.

For artificial infection, three isolates of *F. culmorum,* Fc002, Fc03, and Fc06 from the fungal culture collection of the Chair of Phytopathology, Technical University of Munich (Hofer et al. 2016; Hoheneder et al. 2022; Linkmeyer et al. 2013) were used. The fungi were grown on potato dextrose agar at 21 °C, 60% relative humidity, and a 12 h cycle of UV plus white light and darkness. Fungal inoculum in form of a spore solution was prepared in autoclaved tap water containing 1 ml/l Tween 80® and was adjusted to a spore concentration of 50,000 conidia/ml containing equal amounts of spores from each strain. Mock treatment consisted of autoclaved tap water and 1 ml/l Tween 80.

Barley spikes were sprayed with either *F. culmorum* spore solution or Mock solution until run-off around mid-flowering (GS 65) and afterwards covered and sealed in plastic bags for 2 days to ensure 99% relative air humidity. Treated spikes were sampled 4 and 7 days post inoculation and immediately frozen in liquid nitrogen. For each cultivar, treatment, and time point, four replicates were collected, each consisting of three pooled spikes.

### DNA extraction

Spike tissue was ground in liquid nitrogen and extraction of genomic DNA from 150 mg samples was performed according the method of *(Fraaije et al. 1999)* with modifications as described by *(Hofer et al. 2016)*.

### Quantification of fungal DNA

The fungal DNA content in immature spike tissue in relation to genomic barley DNA was determined via qPCR using species specific primers targeting elongation factor 1α (EF1α) (Nicolaisen et al. 2009). *Fusarium culmorum: FculC561 forward, CACCGTCATTGGTATGTTGTCACT; Fcul614 reverse, CGGGAGCGTCTGATAGTCG*. *Hordeum vulgare*: Hor1 forward, TCTCTGGGTTTGAGGGTGAC; Hor2 reverse, GGCCCTTGTACCAGTCAAGGT. As standards a 10-fold dilution series of target DNA was used. Measurements were performed in two technical replicates using Takyon Low ROX SYBR 2X MasterMix Blue dTTP (Eurogentec, Belgium) with an AriaMx real-time PCR system (Agilent Technologies).

### RNA extraction

Spike material was ground in liquid nitrogen and extraction of total RNA was performed with 100 mg of each sample using the GeneMATRIX Universal RNA Purification Kit (EURx Molecular Biology Products) following the manufacturer’s instructions.

### 3’-RNA Sequencing

Libraries for 3’-RNA-sequencing (Moll et al. 2014) were prepared using the QuantSeq 3’mRNA-Seq Library Prep Kit (Lexogen, Austria) according to the manufactureŕs instructions. Library sizes were quantified with a Qubit Fluorometer 2.0 and a Qubit DNA High Sensitivity Assay Kit, ready-to-load pools of cDNA libraries were determined with qPCR. An Agilent 2010 Electrophoresis Bioanalyzer with the High Sensitivity DNA Assay was used to check the integrity and mRNA fragment size distribution of each library. Next-Generation Sequencing was performed on an Illumina NovaSeq 6000 system.

### 3’RNAseq read quality check, mapping and annotation

The obtained 3’RNAseq reads were processed with nf-core/rnaseq (v3.17.0) using Nextflow (v24.04.4) (Ewels et al. 2020). For mapping we used the published reference genome Morex V3 (Mascher et al. 2021) merged with the *F. culmorum* genome FusCulm_01 (submitted GenBank assembly GCA_003033665.1), with the 3’-UTR regions of the last exon extended either by 3 kb or until the following gene. STAR (v2.7.11b) was used for read alignment, Salmon (v1.10.3) for transcript quantification, FastQC (v0.12.1) for raw read quality control, and Trim Galore (v0.6.10) with Cutadapt (v4.9) for adapter trimming and filtering. The raw reads and mapped data are deposited to GEO and are available under project accession GSE310570 and can be excessed for review using the token mhqbuskkxpaxlmj.

### Differential gene expression analysis

For the differential gene expression analyses, we used the edgeR package version 4.0.16 (Robinson et al. 2010). The mapped *Fusarium* genes were excluded from any downstream analysis. Genes with low expression were filtered by retaining only those with CPM > 10 in at least 4 samples. The dataset was then normalized using TMM normalization via calcNormFactors(). A design matrix was constructed with model.matrix() to define sample groupings. Dispersion estimates were calculated using estimateDisp() with robust estimation. A generalized linear model (GLM) was fitted using glmFit(), and specific contrasts between conditions and genotypes were defined using makeContrasts(genotype_infected_4dpi –genotype_mock_4dpi; genotype_infected_7dpi – genotype_mock_7dpi) to enable differential expression analysis.

### Protein extraction and sample preparation

Proteins were extracted from 500 mg of each sample using trichloracetic acid/acetone precipitation followed by phenolic extraction as described by (Brajkovic et al. 2023). Proteins were quantified via bicinchoninic acid assay (BCA, Thermo Pierce). Tryptic Digest of 200 µg protein lysate was performed following protein aggregation capture on a Bravo Agilent pipetting system using Sera-Mag^TM^ Carboxylate-Modified Magnetic Beads (Cytiva Europe GmbH, Freiburg im Breisgau, Germany).

For Quantitative proteomic measurements, samples were split into 8 batches, consisting of samples from one genotype either 4 or 7 days post inoculation, respectively. Each batch contained 8 samples and one Reference channel with a pooled sample as an inter-run calibrator. Digested Peptides were labelled using 0.1 mg TMT10plex™ Isobaric Mass Tag Labeling Reagent (Thermo Scientific™) for each sample and were pooled according to the batch design. Desalting of labelled and pooled samples was done using C_18_ Sep-Pak cartridges (37–55 µm particle size, 50 mg capacity, Waters) and washing steps were performed with 0.1% formic acid. Peptides were eluted in 0.1% formic acid in 60% Acetonitrile. Eluted peptides were freeze-dried and stored at −20 °C.

### Offline basic reversed-phase fractionation

Basic reversed-phase (bRP) fractionation of peptides was performed as described by (Höfer et al. 2025). The dried peptides were dissolved in 200 µl 25 mM ammonium bicarbonate (pH 8.0). A Vanquish HPLC (Thermo Scientific) was then used for fractionation. The sample was injected onto a Waters BEH130 XBridge C18 column (3.5 µm, 4.6 × 250 mm). Peptides were eluted using a 60 min gradient ranging from 7% to 45% acetonitrile in the constant presence of 2.5 mM ammonium bicarbonate (pH 8.0) at a flow rate of 1000 µl/min. Between 7 and 55 minutes after injection fractions were collected with an automated fraction collector. The resulting 96 fractions were acidified with 1% formic acid and pooled to 48 fractions, before they were freeze-dried and stored at −20 °C.

### LC-MS/MS Measurement

Full proteome-TMT peptides were measured with an Eclipse Tribrid mass spectrometer (Thermo Scientific) that was coupled to a Vanquish Neo (Thermo Scientific). The sample was directly injected onto the Acclaim PepMap 100 C18 column (2 µm particle size, 1 mm ID × 150 mm). Separation was performed on a 27 min segmented gradient with a flow rate of 50 µl/min starting from 4% to 27% B (23min) and 27% to 32% B (2min). The system was finally washed with 100 μl 90% B and re-equilibrated at 1% B. Solvent A consisted of 0.1 %v FA and 3 %v DMSO in water. Solvent B consisted of 0.1%v FA and 3%v DMSO in Acetonitrile. The MS was operated in a fast, data-dependent MS3 - mode. The spray voltage was set to 3.5 kV supported by sheath gas (32 units) and aux gas (5 units) with a vaporizer temperature of 125 °C. Every 1.2 s, a full-scan (MS1) was recorded from 360 to 1600 m/z with a resolution of 60k in the Orbitrap in profile mode. The MS1 AGC target was set to 4e5. Based on the full scans, precursors were targeted for MSMS scans if the charge was between 2 and 6, the isotope envelop was peptidic (MIPS), and the intensity exceeded 1e4. The MS2 quadrupole isolation window was set to 0.6 Th. The TMT peptides were HCD fragmented with an NCE of 34%. The MS2 spectra were acquired in the ion trap in rapid mode. The MS2 AGC target was set to 3e4 charges, and the maxIT was set to 40 ms. The maxIT or AGC target could be dynamically exceeded when the previous scan took longer than the calculated injection time (inject beyond mode). Precursors that have been targeted for fragmentation were excluded for 50 s for all possible charge stages. TMT reporter ions were measured in a consecutive MS3 scan based on the previous MSMS scan. Thus, a new batch of precursor ions was isolated with an MS3 quadrupole isolation window of 1.2. The isolated precursor was then HCD-fragmented identically to the previous MS2 scan. Additionally, Isobaric tag loss exclusion properties were set to TMT reagent. The selected fragment ions were then HCD fragmented with an NCE of 55%. The MS3 spectrum was acquired with 50k resolution from 100 to 1000 Th in the Orbitrap in centroid mode. The MS3 AGC target was set to 2e5 charges, and the maxIT was set to 86 ms.

### Protein Identification

The mass-spectrometry data was processed using MaxQuant (version 2.4.2.0) for peptide and protein identification and quantification. The measured spectra were searched against *Hordeum vulgare* Morex v3 genome-based protein prediction (Mascher et al. 2021) and *Fusarium culmorum* UniProt reference proteome UP000241587 (The UniProt Consortium 2023). The default MaxQuant parameters were used except for enzyme specificity set to Trypsin/P, allowing for two missed cleavages. Carbamidomethylation at cysteine residues was defined as a fixed modification, while N-terminal acetylation and oxidation of methionine were set to variable. TMT-based quantification was performed based on reporter ion intensities. The false discovery rate (FDR) at peptide spectrum match (PSM) level was fixed to 1%.

The raw data is deposited to PRIDE under project accession PXD068736 and can be excessed for review using the following details: Username; Password: JkCNkrHM5JSmp.

### Differential Protein Expression Analysis

Reporter intensities were normalized for further analysis, using a twostep procedure. First, sample-wise normalization was performed to account for differences in total protein loading. A sample specific normalization factor was calculated by dividing the mean sum intensity across all samples by the summed intensity of each individual sample. Raw intensity values were then multiplied by their respective correction factor. In a second step, values were corrected for batch effects using protein-specific correction factors calculated using the median intensity of a protein in the reference channels divided by the protein intensity measured in the reference channel in each batch. The batch-specific correction factor was then multiplied with the sample intensities in the corresponding batch. Normalization was carried out in R using **tidyverse (Wickham et al. 2019).**

Normalized protein intensities were processed in Perseus (version 2.0.11). Contaminants, Reverse hits and proteins mapped to *Fusarium culmorum* were removed from the dataset. Additionally, the data was filtered to only include proteins with at least one valid value across all samples. The samples were grouped according to barley genotype, sampling time point and treatment. Data was log_2_ transformed and filtered again to retain proteins containing at least 3 valid values in at least one experimental group. Missing values were then imputed based on normal distribution (width: 0.3; down shift: 1.8).

The data was analyzed in Perseus (version 2.0.11), performing a principal component analysis (PCA), hierarchical clustering as well as a differential expression analysis (Student’s *t*-test) comparing Mock treated to *F. culmorum* infected samples, assuming significance at p < 0.05.

### GO-term Enrichment Analysis

GO-term enrichment analysis of the DEGs and DEPs was performed using the topGO R-package v.2.52.0 (http://dx.doi.org/10.18129/B9.bioc.topGO). GO-term annotation for barley genes were downloaded from G:Profiler at https://biit.cs.ut.ee/gprofiler/gost (Kolberg et al. 2023). As the gene universe for analyzing DEGs the transcripts with a threshold CPM>1 in at least 1 sample (36920) were used. The gene universe for the analysis of DEPs consisted of all quantified proteins (7671). Statistical significance of the enrichment was calculated using Fisher’s exact test. The enrichment was considered significant for p-values < 0.01.

### KEGG Pathway Analysis

Functional annotation of barley was performed using BlastKOALA (https://www.kegg.jp/blastkoala/) by performing a BLAST search of the Morex V3 genome against KEGG annotated eukaryotic genomes. This search assigned KEGG onthology (KO) numbers to 30% of all barley gene IDs (Supplement Table D1.6).

KEGG Pathway Mapping was done using KEGG Mapper’s Reconstruct function (https://www.kegg.jp/kegg/mapper/reconstruct.html) to match differentially regulated genes and proteins with metabolic functions. Enriched pathway modules were identified and used to reconstruct relevant metabolic pathways.

### Untargeted Metabolomics Analysis

For untargeted metabolomics analysis of *F. culmorum*-infected barley spikes, 100 mg of frozen plant material was weighed into 2-ml bead-beater tubes (Bertin Technologies) filled with zirconium oxide beads (1.4 and 2.8 mm), and homogenized in 1 ml methanol/water (70/30, v/v) using a Precellys® homogenizer at 6500 rpm for 3 × 30 s with 15 s pauses. After centrifugation, the supernatant was filtered (0.45 µm, Sartorius) and used for UHPLC-TOF-MS analysis.

Secondary metabolite analysis was performed via UHPLC on an ACQUITY I-Class system with a BEH C18 column (150 × 2.1 mm, 1.7 µm, Waters), using 0.1% formic acid in water (eluent A) and acetonitrile (eluent B). The gradient was 1% B (1 min), 1–35% B (6 min), 35–70% B (1 min), 70–100% B (1 min), hold at 100% B (3 min), 100–1% B (0.5 min), and 1% B (1 min), with a flow rate of 0.4 ml/min, autosampler at 10 °C, column at 45 °C, and injection volume of 3 µl.

Mass spectrometry was performed using a Synapt G2-S HDMS mass spectrometer (Waters, Manchester, UK) in high-resolution mode and electrospray ionization (ESI) with a scan time for the MS^E^ method (centroid) of 0.1 s. The instrument was operated in positive and negative ion mode, applying the following source parameters: capillary voltage +2.5 kV (ESI^+^), –3.0 kV (ESI^−^), sampling cone 20 V, source offset 40 V, source temperature 120 °C, desolvation temperature 450 °C, cone gas flow 2 L/h, nebulizer 6.5 bar and desolvation gas 850 L/h, and collision energy ramp 20–40 eV. All data were lock mass corrected on the pentapeptide leucine enkephaline (Tyr-Gly-Gly-Phe-Leu, *m/z* 554.2615 [M-H]^−^) in a solution (1 ng/µL) of acetonitrile/0.1% formic acid 1/1 (v/v). Scan time for the lock mass was set to 0.3 s with an interval of 15 s, and three scans were made on average with a mass window of ±0.3 Da. Calibration of the Synapt G2-S in the range of *m/z* 50 to 1,200 was performed using a solution of sodium formate (5 mmol/L) in 2-propanol/water 9/1 (v/v).

The raw files are deposited to zenodo and are available under https://doi.org/10.5281/zenodo.18391760.

Mass spectrometry data were analyzed using MassLynx software (version 4.1 SCN 901, Waters, Manchester, UK). Principal component analysis (PCA) for untargeted metabolomics and lipidomics was conducted using Progenesis QI software (version 3.0, Waters, Manchester, UK) applying the following peak picking conditions: all runs, limits automatic, sensitivity 3, and retention time limits 0.5–11.5 min. Metabolic feature annotation was performed in Progenesis QI using a custom compound database generated from curated molfiles that were imported to enable accurate matching of detected features based on exact mass and retention time. Metabolic feature identification was assigned on three levels: confirmed IDs based on reference standards measured under identical analytical conditions (ID-level 1), putative annotations derived from physicochemical properties matched to the custom database and public libraries (ID-level 2), and putative compound-class assignments based on characteristic fragmentation patterns (ID-level 3).

Compounds used for PCA were filtered by means of ANOVA *p* ≤ 0.05 and fold change ≥ 2. The processed data were exported to EZinfo (version 3.0, Waters, Manchester, UK), and the matrix was analyzed by PCA with pareto scaling. Group differences were calculated using orthogonal partial least squares discriminant analysis (OPLS-DA) and visualized in S-plots.

## ETHICS APPROVAL AND CONSENT TO PARTICIPATE

Not applicable

## CONSENT FOR PUBLICATION

Not applicable

## AVAILABILITY OF DATA AND MATERIAL

The raw 3’RNA-sequencing reads and processed alignment files have been deposited in the NCBI Gene Expression Omnibus (GEO) under accession number GSE310570. The data can be accessed for review using the secure token *mhqbuskkxpaxlmj*.

The raw proteomics data have been deposited in the PRIDE repository under the project accession PXD068736. The dataset is available for review using the following credentials: **Username**; **Password**: JkCNkrHM5JSm

Raw metabolomic data are deposited on zenodo and are available under https://doi.org/10.5281/zenodo.18391760.

## AUTHORS’ CONTRIBUTIONS

S.H., C.E.S., F.H., and R.H. designed the experiments; S.H., C.E.S., and F.H. performed the greenhouse experiments and sample preparations; Proteomics measurements were performed by S.B. in the lab of B.K.; Metabolomics measurements were performed by L.K. in the lab of C.D.; The data was analyzed by S.H., C.E.S., L.K., and T.D.S.; S.H. wrote the manuscript supported by R.H., with contribution from S.B. and C.E.S.;. R.H., C.D. and B.K. acquired funding. All authors read and approved the final manuscript.

## ACKNOWLEDGMENTS

We thank Verena Klingl for her support in sample preparation and Regina Dittebrandt for the cultivation of fungal strains. We thank the staff at the Greenhouses & Phytochambers Unit (GPU) at the Plant Technology Center of the Technical University of Munich for careful barley cultivation. We thank Dr. Christine Wurmser (Division of Animal Physiology and Immunology, TUM School of Life Sciences, Technical University of Munich, Freising, Germany) for Library preparation and organization of 3’RNA sequencing. We thank Ingo Fritz (Computational Neurosciences, TU Munich) for mapping the 3’RNAseq data.

## Supplemental Figures

**Fig. S1:**
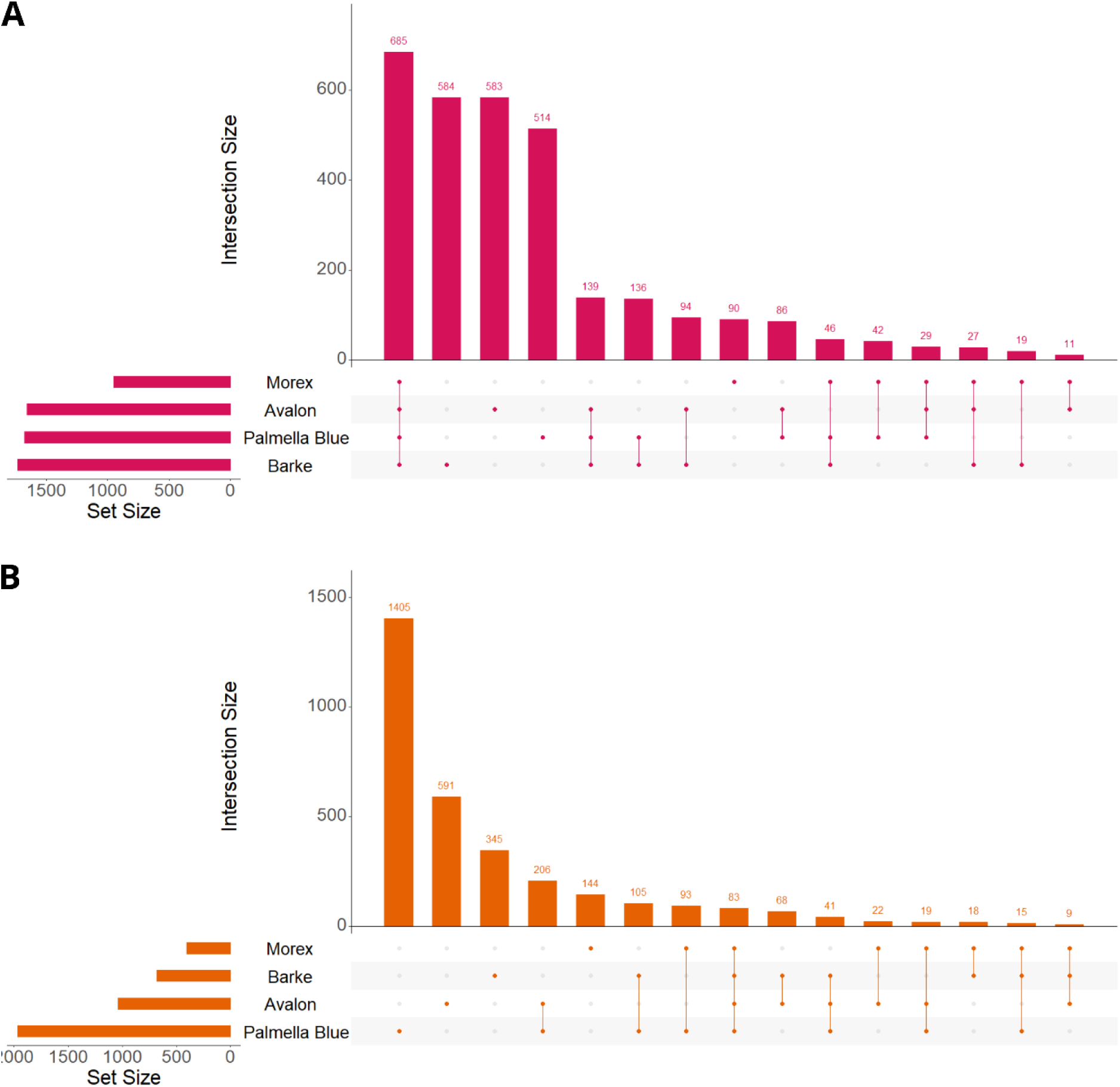
UpSet plot illustrating the intersections among the DEGs (**A**) and DEPs (**B**) identified in four barley cultivars Avalon, Barke, Morex and Palmella Blue. Horizontal bars on the left represent the total number of DEPs or DEGs in each individual cultivar. The matrix indicates which sets participate in each intersection, with connected dots marking a given combination. Vertical bars above the matrix show the size of each intersection.

**Fig. S2.**
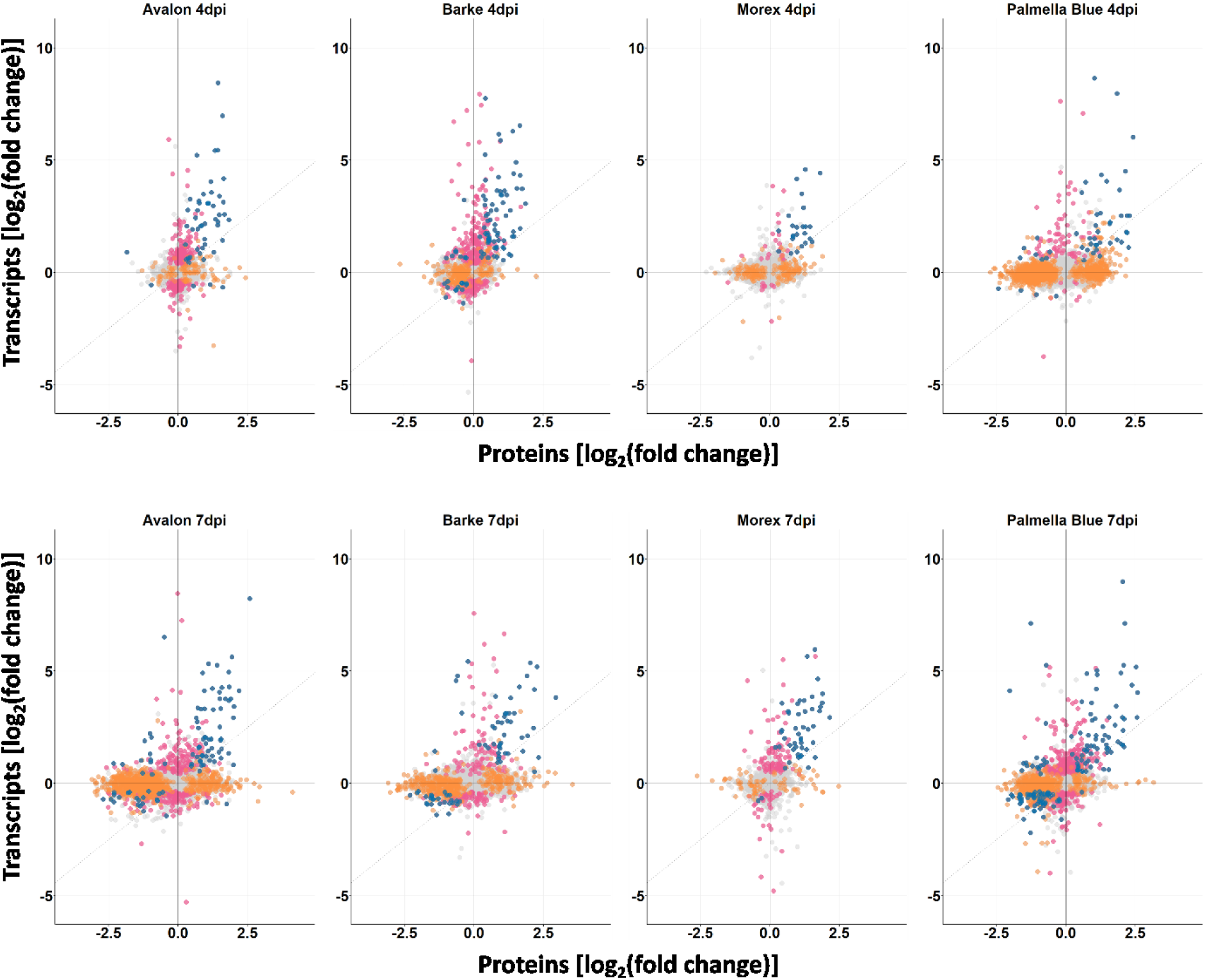
Comparison of fold changes of transcripts and corresponding proteins in barley cultivars Avalon, Barke, Morex, and Palmella Blue at 4 and 7 days post inoculation. Pink: Differentially expressed genes (DEGs). Orange: Differentially abundant proteins (DEPs). Blue: DEP/DEG pairs. Inoculation was performed around mid-anthesis. Infected samples were compared with mock-treated controls of the same cultivar and time-point. For each cultivar, treatment, and time point, four replicates were collected, each consisting of three pooled spikes.

**Fig. S3.**
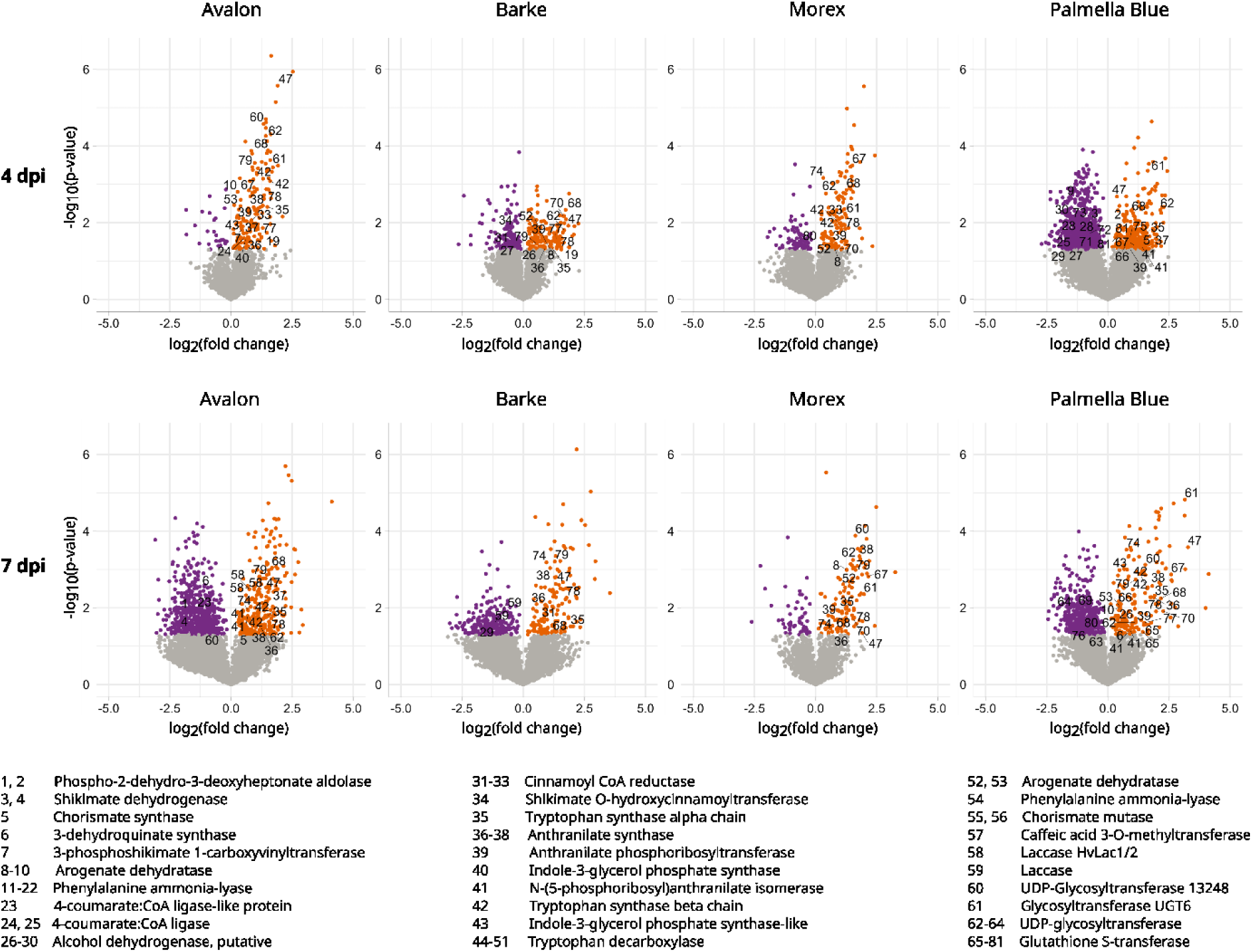
Volcano plots of the differential abundance of proteins in barley cultivars Avalon, Barke, Morex, and Palmella Blue at four and seven days post inoculation with *F. culmorum* spore solution in comparison with mock-treated controls. Inoculation was performed around mid-anthesis. Infected samples were compared with mock-treated controls of the same cultivar and time-point. For each cultivar, treatment, and time point, four replicates were collected, each consisting of three pooled spikes. Volcano plots show the log_2_(fold change) values in relation to the negative log_10_(p-value).

**Fig. S4.**
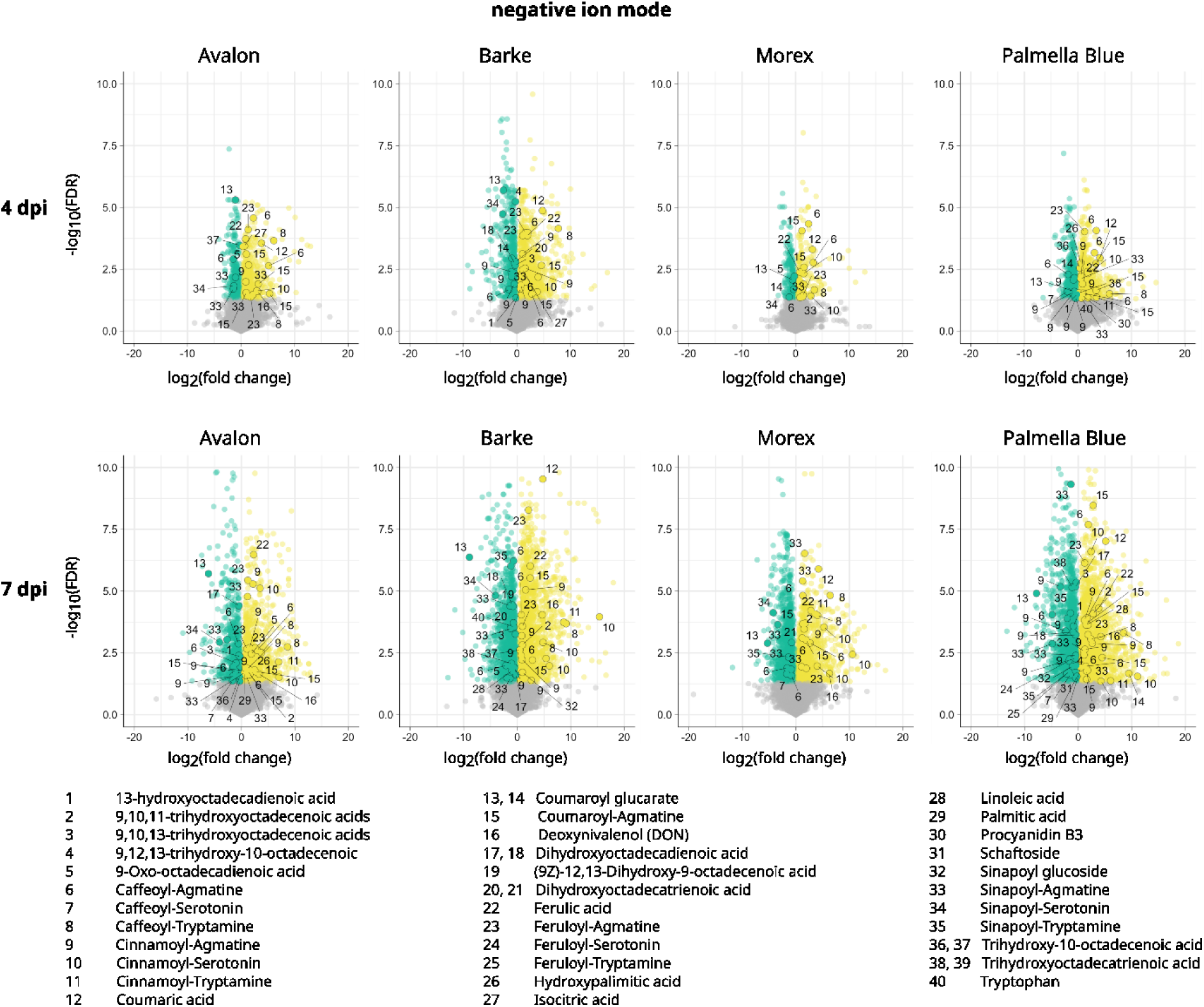
Volcano plots of the differential abundance of metabolic features in barley cultivars Avalon, Barke, Morex, and Palmella Blue at four and seven days post inoculation with *F. culmorum* spore solution in comparison with mock-treated controls. Metabolic features were measured via mass spectrometry in negative ion mode. Inoculation was performed around mid-anthesis. Infected samples were compared with mock-treated controls of the same cultivar and time-point. For each cultivar, treatment, and time point, four replicates were collected, each consisting of three pooled spikes. Volcano plots show the log_2_(fold change) values in relation to the negative log_10_(FDR).

**Fig. S5.**
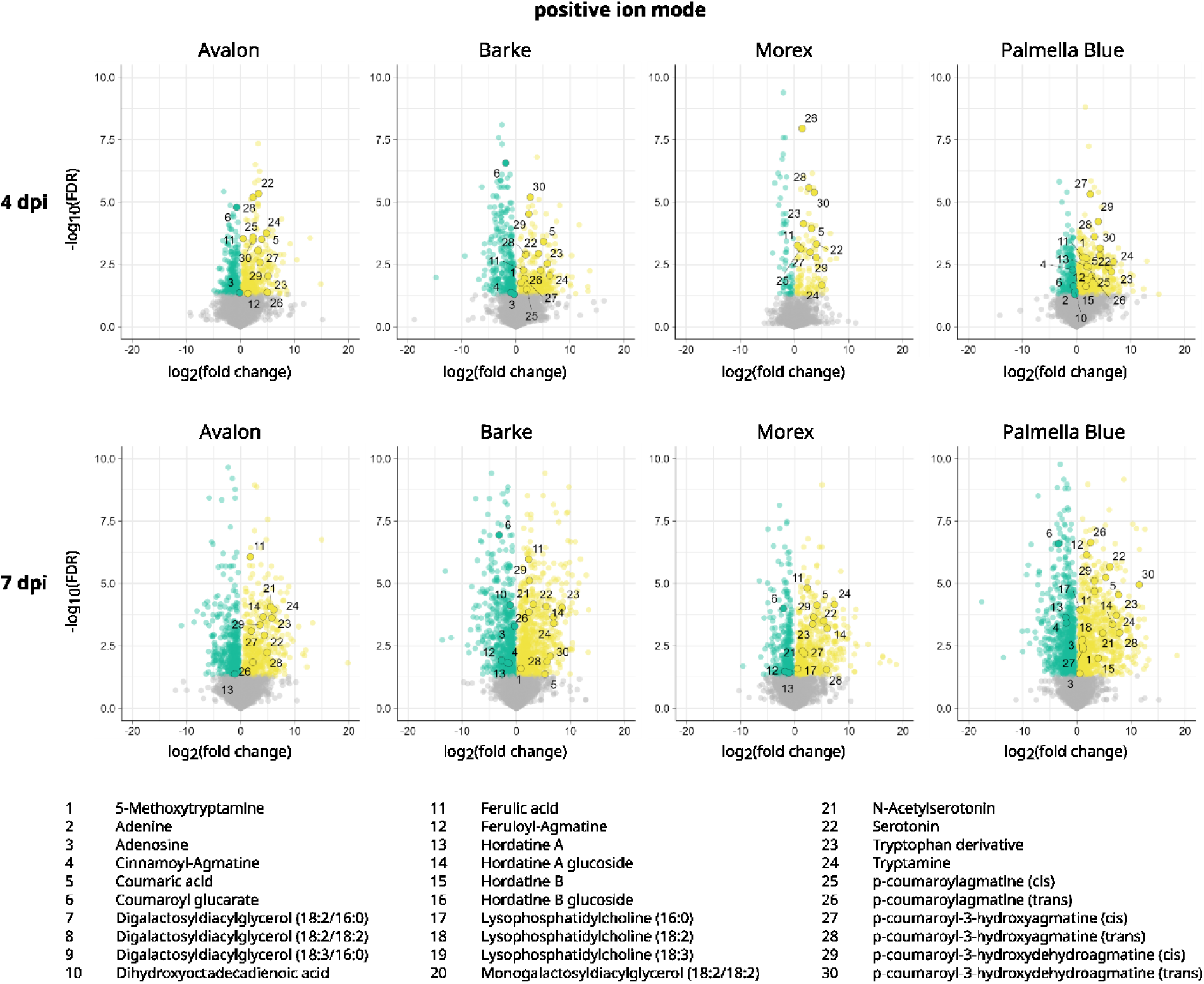
Volcano plots of the differential abundance of metabolic features in barley cultivars Avalon, Barke, Morex, and Palmella Blue at four and seven days post inoculation with *F. culmorum* spore solution in comparison with mock-treated controls. Metabolic features were measured via mass spectrometry in positive ion mode. Inoculation was performed around mid-anthesis. Infected samples were compared with mock-treated controls of the same cultivar and time-point. For each cultivar, treatment, and time point, four replicates were collected, each consisting of three pooled spikes. Volcano plots show the log_2_(fold change) values in relation to the negative log_10_(FDR).

**Fig. S6.**
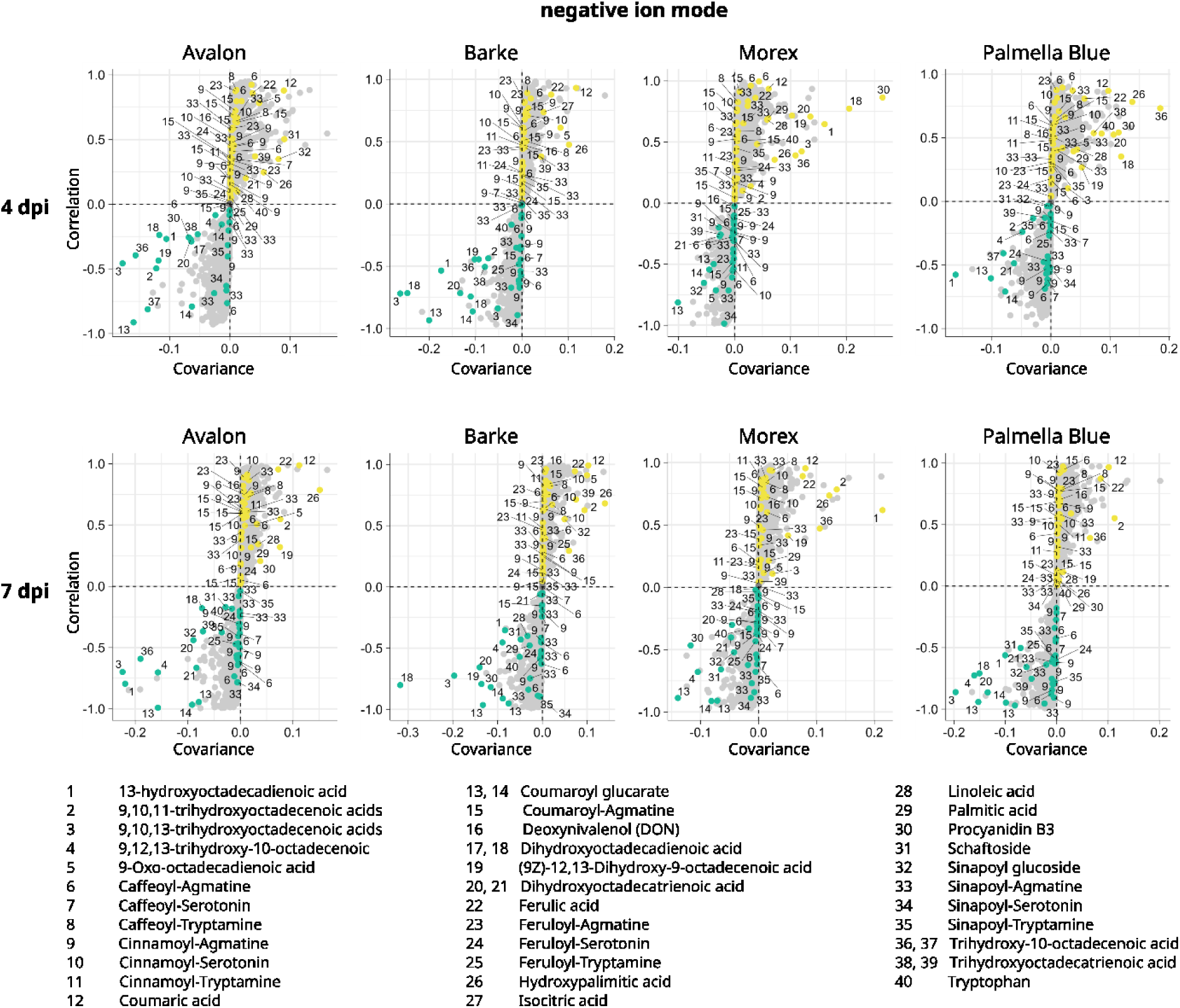
S-plots of the differential abundance of metabolic features in barley cultivars Avalon, Barke, Morex, and Palmella Blue at four and seven days post inoculation with *F. culmorum* spore solution in comparison with mock-treated controls. Metabolic features were measured via mass spectrometry in negative ion mode. Inoculation was performed around mid-anthesis. Infected samples were compared with mock-treated controls of the same cultivar and time-point. For each cultivar, treatment, and time point, four replicates were collected, each consisting of three pooled spikes. S-plots show group differences as calculated via orthogonal partial least squares discriminant analysis (OPLS-DA).

**Fig. S7.**
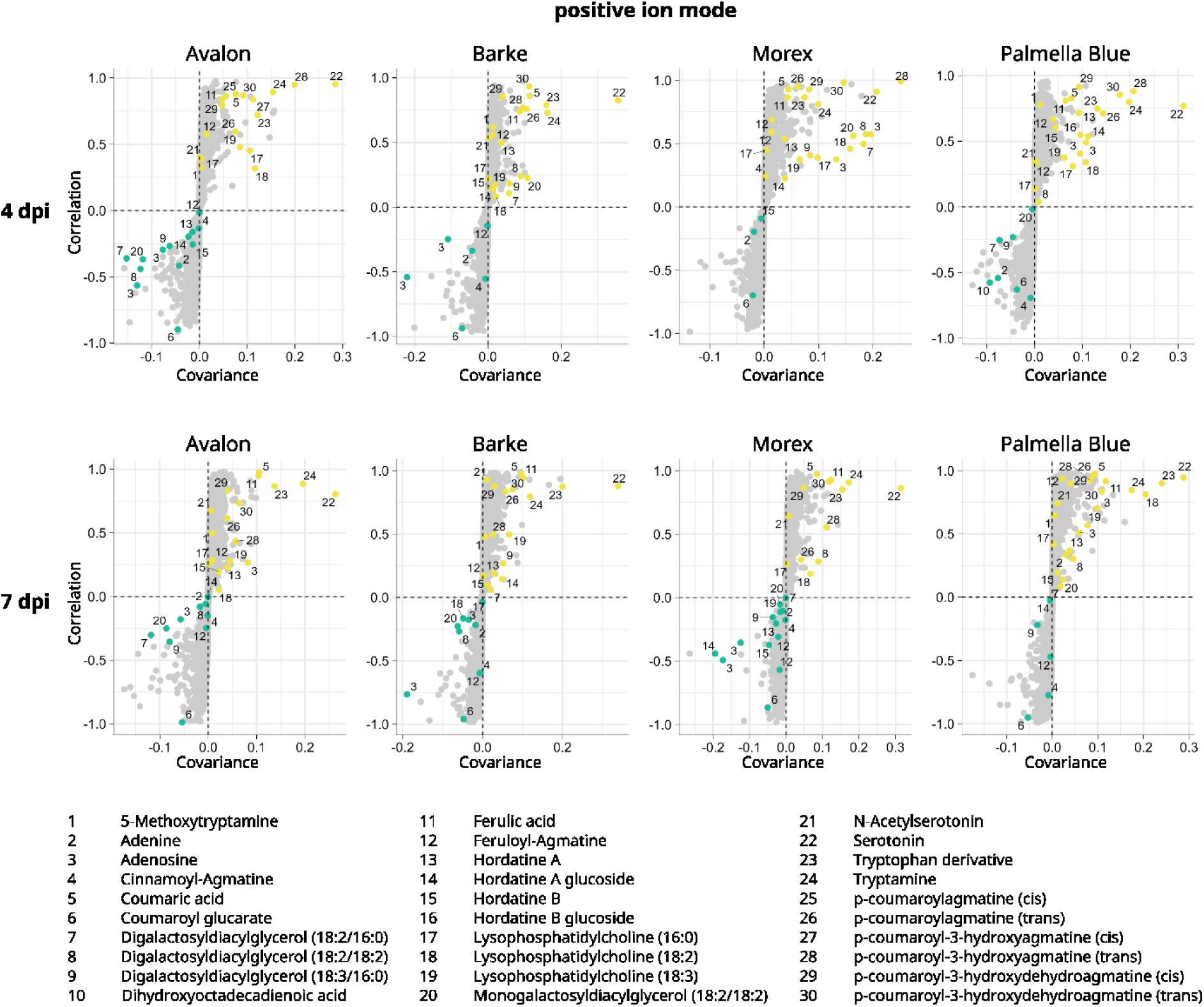
S-plots of the differential abundance of metabolic features in barley cultivars Avalon, Barke, Morex, and Palmella Blue at four and seven days post inoculation with *F. culmorum* spore solution in comparison with mock-treated controls. Metabolic features were measured via mass spectrometry in positive ion mode. Inoculation was performed around mid-anthesis. Infected samples were compared with mock-treated controls of the same cultivar and time-point. For each cultivar, treatment, and time point, four replicates were collected, each consisting of three pooled spikes. S-plots show group differences as calculated via orthogonal partial least squares discriminant analysis (OPLS-DA).

